# Overview of the SAMPL6 host-guest binding affinity prediction challenge

**DOI:** 10.1101/371724

**Authors:** Andrea Rizzi, Steven Murkli, John N. McNeill, Wei Yao, Matthew Sullivan, Michael K. Gilson, Michael W. Chiu, Lyle Isaacs, Bruce C. Gibb, David L. Mobley, John D. Chodera

## Abstract

Accurately predicting the binding affinities of small organic molecules to biological macro-molecules can greatly accelerate drug discovery by reducing the number of compounds that must be synthesized to realize desired potency and selectivity goals. Unfortunately, the process of assessing the accuracy of current computational approaches to affinity prediction against binding data to biological macro-molecules is frustrated by several challenges, such as slow conformational dynamics, multiple titratable groups, and the lack of high-quality blinded datasets. Over the last several SAMPL blind challenge exercises, host-guest systems have emerged as a practical and effective way to circumvent these challenges in assessing the predictive performance of current-generation quantitative modeling tools, while still providing systems capable of possessing tight binding affinities. Here, we present an overview of the SAMPL6 host-guest binding affinity prediction challenge, which featured three supramolecular hosts: octa-acid (OA), the closely related tetra-endo-methyl-octa-acid (TEMOA), and cucurbit[8]uril (CB8), along with 21 small organic guest molecules. A total of 119 entries were received from 10 participating groups employing a variety of methods that spanned from electronic structure and movable type calculations in implicit solvent to alchemical and potential of mean force strategies using empirical force fields with explicit solvent models. While empirical models tended to obtain better performance than first-principle methods, it was not possible to identify a single approach that consistently provided superior results across all host-guest systems and statistical metrics. Moreover, the accuracy of the methodologies generally displayed a substantial dependence on the system considered, emphasizing the need for host diversity in blind evaluations. Several entries exploited previous experimental measurements of similar host-guest systems in an effort to improve their physical-based predictions via some manner of rudimentary machine learning; while this strategy succeeded in reducing systematic errors, it did not correspond to an improvement in statistical correlation. Comparison to previous rounds of the host-guest binding free energy challenge highlights an overall improvement in the correlation obtained by the affinity predictions for OA and TEMOA systems, but a surprising lack of improvement regarding root mean square error over the past several challenge rounds. The data suggests that further refinement of force field parameters, as well as improved treatment of chemical effects (e.g., buffer salt conditions, protonation states) may be required to further enhance predictive accuracy.

## Introduction

Quantitative physical and empirical modeling approaches have played a growing role in aiding and directing the design of small molecule biomolecular ligands for use as potential therapeutics or chemical probes [1 - 4, 23, 65]. The degree of inaccuracy of these predictions largely determines how effective they can be in prioritizing synthesis of small molecule ligands [105]. Retrospective estimates have suggested that current methodologies are capable of achieving about 1-2 kcal/mol inaccuracy for well-behaved protein-ligand systems [6, 122], but more work remains to be done to extend the applicability domain of these technologies.

Assessment of how much of this inaccuracy can be attributed to fundamental limitations of the *force field* in accurately modeling energetics is complicated by the presence of numerous additional factors [78]. Proteins are highly dynamic entities, and many common drug targets—such as kinases [115] and GPCRs [63]—possess slow dynamics with timescales of microseconds to milliseconds [62] that frustrate the computation of true equilibrium affinities. While there has been some attempt to curate benchmark sets of protein-ligand affinity data in well-behaved model protein-ligand systems that are believed to be mostly free of slow-timescale motions that would convolve convergence issues with forcefield inaccuracies [78], other effects can complicate assessment of the accuracy of physical modeling benchmarks. Ionizable residues, for example, comprise approximately 29% of all protein residues [56], and large-scale computational surveys suggest that 60% of all protein-ligand complexes undergo a change in ionization state upon binding [5], with several notable cases characterized experimentally [24, 25, 91, 111]. For physical or empirical modeling approaches that assume fixed protonation states throughout the complexation process, protonation state effects are hopelessly convolved with issues of force field inaccuracy.

### Host-guest systems are a tractable model for assessing force field inaccuracies

Over the last decade, supramolecular host-guest complexes have emerged as a practical and useful model system for the quantitative assessment of modeling errors for the interaction of druglike small molecules with receptors. Supramolecular hosts such as cucurbiturils, cavitands, and cyclodextrins can bind small druglike molecules with affinities similar to protein-ligand complexes [84, 85, 99]. The lack of slowly relaxing conformational degrees of freedom of these hosts eliminates the potential for slow microsecond-to-millisecond receptor relaxation timescales as a source of convergence issues [78], while the small size of these systems allows many methodologies to take advantage of faster simulation times to rapidly assess force field quality. The high solubilities of these systems permit high-quality biophysical characterization of their interactions via gold-standard methods such as isothermal titration calorimetry (ITC) and nuclear magnetic resonance (NMR) [22, 38, 114]. Additionally, the stability of supramolecular hosts at extreme pH allows for strict control of protonation states in a manner not possible with protein-ligand systems, allowing confounding protonation state effects to be eliminated from consideration if desired [114]. Collectively, these properties have made host-guest systems a productive route for revealing deficiencies in modern force fields through blind community challenge exercises we have organized as part of the Statistical Assessment of the Modeling of Proteins and Ligands (SAMPL) series of blind prediction challenge [86, 88, 107, 127].

### SAMPL host-guest challenges have driven advances in our understanding of sources of error

The SAMPL (Statistical Assessment of the Modeling of Proteins and Ligands) challenges are a recurring series of blind prediction challenges for the computational chemistry community [76, Drug Design Data Resource]. Through these challenges, SAMPL aims to evaluate and advance computational tools for rational drug design: By focusing the community on specific phenomena relevant to drug discovery—such as the contribution of force field inaccuracy to binding affinity prediction failures—isolating these phenomena from other confounding factors in well-designed test systems, evaluating tools prospectively, enforcing data sharing to learn from failures, and releasing the resulting high-quality datasets into the community as benchmark sets, SAMPL has driven progress in a number of areas over five previous rounds of challenge cycles [9, 34, 35, 43, 44, 81, 82, 86, 88, 93, 107, 107, 108, 127].

More specif cally, SAMPL host-guest challenges have provided key tests for modeling of binding interactions [78], motivating increased attention to how co-solvents and ions modulate binding (resulting in errors of up to 5 kcal/mol when these effects are neglected) and the importance of adequately sampling water rearrangements [17, 78, 86, 127]. In turn, this detailed examination has resulted in clear improvements in subsequent SAMPL challenges [127], though host-guest binding remains difficult to model accurately [47], in part due to force field limitations (spawning new efforts to remedy major force field defficiencies [125]).

### SAMPL6 host-guest systems

Three hosts were selected for the SAMPL6 host-guest binding challenge from the Gibb Deep Cavity Cavitand (GDCC) [36, 48, 79, 80] and the cucurbituril (CB) [31, 69, 83] families (*Figure 1*). The guest ligand sets were purposefully selected for the SAMPL6 challenge. The utility of these particular host systems for evaluating free energy calculations has been reviewed in detail elsewhere [79, 80].

**Figure 1.**
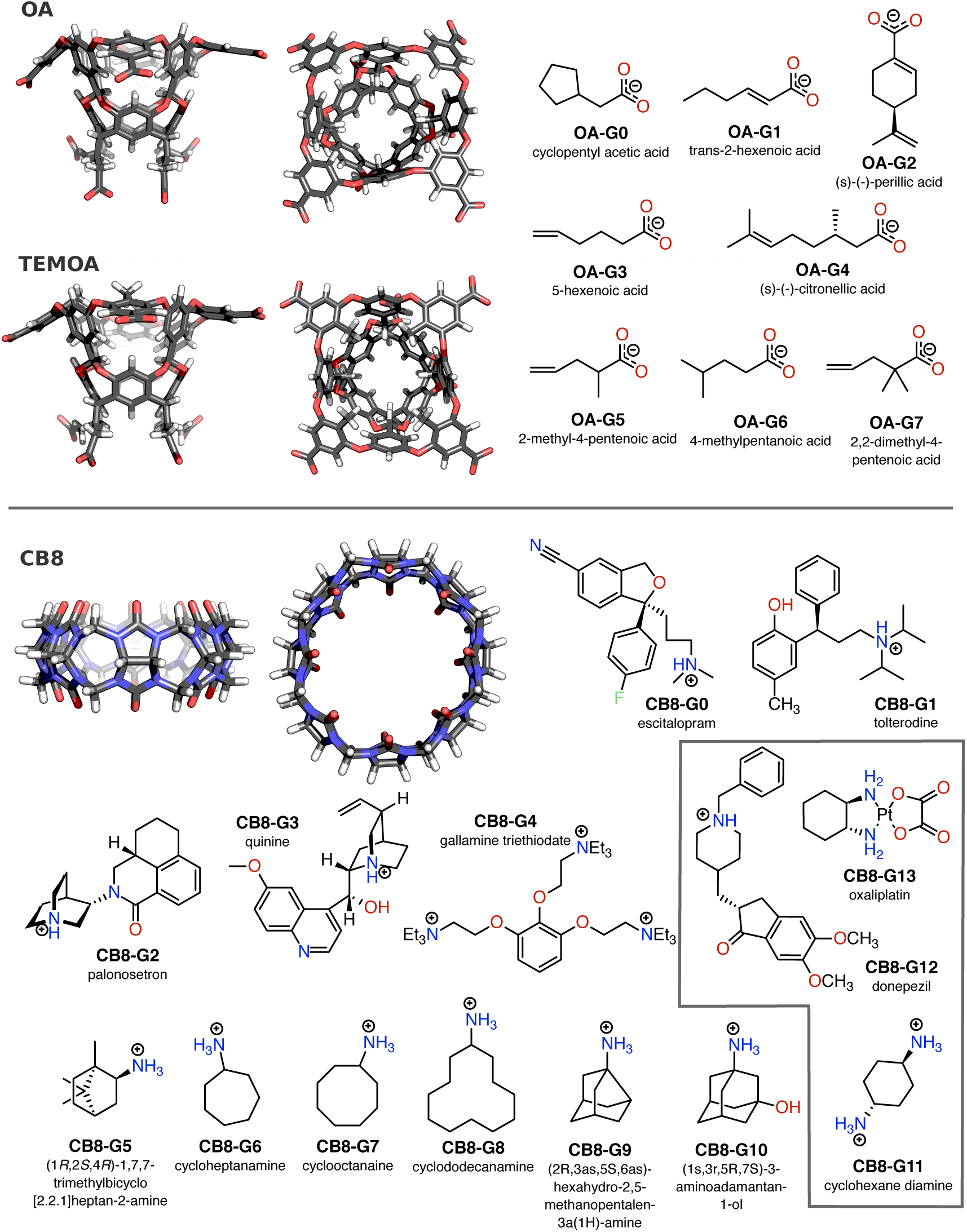
Hosts and guests featured in the SAMPL6 host-guest blind challenge dataset. Three-dimensional structures of the three hosts featured in the SAMPL6 challenge dataset (OA, TEMOA, and CB8) are shown in stick view from top and side perspective views. Carbon atoms are represented in gray, hydrogens in white, nitrogens in blue, and oxygens in red. Guest ligands for each complex are shown as two-dimensional chemical structures annotated by hyphenated host and guest names. Protonation states of the guest structures correspond to the predicted dominant microstate at the experimental pH at which binding affinities were collected, and matches those provided in the `moi2` and `sdf` input files shared with the participants when the challenge was announced. The same set of guests OA-G0 through OA-G7 was used for both OA and TEMOA hosts. The gray frame (lower right) contains the three CB8 guests that constitute the bonus challenge.

The two GDCCs, octa-acid (OA) [36] and tetra-endo-methyl-octa-acid (TEMOA) [33], are low-symmetry hosts with a basket-shaped binding site accessible through the larger entryway located at the top. These hosts also appeared in two previous SAMPL host-guest challenges—SAMPL4 [86] and SAMPL5 [127]—with the names of OAH and OAMe respectively with different sets of guests. OA and TEMOA differ by four methyl groups that reduce the size of the binding site entryway (*Figure 1*). Both hosts expose eight carboxyl groups that increase their solubility. The molecular structures of the eight guests selected for the SAMPL6 challenge for characterization against both OA and TEMOA are shown in *Figure 1* (denoted OA-G0 through OA-G7). These guests feature a single polar group situated at one end of the molecule that tends to be exposed to solvent when complexed, while the rest of the compound remains buried in the hydrophobic binding site.

A second set of guest ligands were developed for the host cucurbit[8]uril (CB8). This host previously appeared in the SAMPL3 host-guest binding challenge [87], but members of the same family or analogs such as cucurbit[7]uril (CB7) and CBClip [128] were featured in SAMPL4 and SAMPL5 challenges as well. CB8 is a symmetric (D_8*h*_), ring-shaped host comprising eight identical glycoluril monomers linked by pairs of methylene bridges. Its top-bottom symmetry means that asymmetric guests have at least two symmetry-equivalent binding modes that can be kinetically separated by timescales not easily achievable by standard molecular dynamics (MD) or Monte Carlo simulations and may require special considerations, in particular in alchemical absolute binding free energy calculations [75]. The CB8 guest set (compounds CB8-G0 to CB8-G13 in *Figure 1*) includes both fragment-like and bulkier drug-like compounds.

Some of the general modeling challenges posed by both families of host-guest systems have been characterized in previous studies. While their relatively rigid structure minimizes convergence difficulties associated with slow receptor conformational dynamics, both families have been shown to bind guest ligands via a dewetting processes—in which waters must be removed from the binding site to accommodate guests—in a manner that can frustrate convergence for strategies based on molecular simulation. In the absence of tight-binding guest ligands, the octa-acid host experiences fluctuations in the number of bound waters on timescales of several nanoseconds [30]; a similar phenomenon was observed in alchemical absolute binding free energy calculations of CB7 at intermediate alchemical states with partially decoupled Lennard-Jones interactions [101]. In addition, hosts in both families have been shown to bind ions that can compete with and lower the binding affinity of other guests in solution [37, 98, 109]. Depending on differences in concentration and composition, the effect on the binding free energy can be between 1–2 kcal/mol [85, 94, 98]. Sensitivity of the guest affinity to ion concentration has been observed also with computational methods [50, 89, 95], which suggests that careful modeling of the buffer conditions is in principle necessary for a meaningful comparison to experiments.

### Experimental host-guest affinity measurements

A detailed description of the experimental methodology used to collect binding affinity data for OA, TEMOA, and CB8 host-guest systems is described elsewhere [90]. Briefly, all host-guest binding affinities were determined via direct or competitive isothermal titration calorimetry (ITC) at 298 K. OA and TEMOA measurements were performed in 10 mM sodium phosphate buffer at pH 11.7±0.1 whereas CB8 guests binding affinities were measured in a 25 mM sodium phosphate buffer at pH 7.4. Phosphate buffer is a common choice of buffer for its relevance to biology, and can be prepared over a wide pH range for exerting control over protonation states. Binding stoichiometries were determined by ^1^H NMR spectral integration and/or by ITC. The ITC titration curves were fitted to a single-site model or a competition model for all guests, except for CB8-G12 (donepezil), for which a sequential binding model was used. The stoichiometry coefficient was either fitted simultaneously with the other parameters or fixed to the value verif ed by the NMR titrations, which is the case for the CB8 guest set, as well as for OA-G5, TEMOA-G5, and TEMOA-G7.

To determine experimental uncertainties, we added the relative error in the nonlinear fit-derived association constant (*K*_*a*_) or binding enthalpy (AH) with the relative error in the titrant concentration in quadrature [20]. We decided to arbitrarily assume a relative error in the titrant concentration of 3% after personal communication with Professor Lyle Isaacs who suggested a value inferior to 5% based on his experience. The minimum relative nonlinear fit-derived uncertainty permitted was 1%, since the ft uncertainty was reported by the ITC software as smaller than this in some cases. It should be noted that the error propagation strategy adopted here assumes that the stoichiometry coefficient is fitted to the ITC data in order to absorb errors in cell volume and titrand concentration; this approach is exact only for the OA/TEMOA sets with the exclusion of OA-G5, TEMOA-G5, and TEMOA-G7, and an underestimate of the true error for the remaining cases. The error was then further propagated to the binding free energies and entropies that were calculated from *K*_*a*_ and Δ*H*. The final estimated experimental uncertainties are relatively small, never exceeding 0.1 kcal/mol.

The resulting experimental measurements with their uncertainties are reported in *Table 1* and *Figure 2*. The dynamic range of the binding free energy Δ*G* spans 4.25 kcal/mol for the merged OA and TEMOA guest set, and 7.05 kcal/mol for CB8. The relatively wide cavity of CB8 enables binding stoichiometries different than 1:1. This is the case for three of the CB8 guests, specif cally CB8-G1 (tolterodine), CB8-G4 (gallamine triethiodate), and CB8-G12 (donepezil). Curiously, while CB8-G12 was found to bind in 2:1 complexes (two guests bound to the same host), the NMR experiments determined stoichiometries of 1:2 and 1:3 for CB8-G1 and CB8-G4 respectively (one guest bound to multiple hosts). For the last two guests, the ITC titration curves ft well to a single set of sites binding model which indicates that the each of the binding events are equivalent. In *Table 1* and *Figure 2* we report the binding affinity of both the 1:1 and the 2:1 complex for CB8-G12, which are identified by CB8-G12a and CB8-G12b respectively, and the free energy of the 1:1 complex for CB8-G1 and CB8-G4.

**Table 1.** Summary of ITC and NMR measurements for the SAMPL6 host-guest dataset. Guest identifiers (ID), association constants (*K*_*a*_), binding free energies (Δ*G*), enthalpies (Δ*H*), entropies at room temperature (*T*Δ*S*) and stoichiometric ratios (*n*) as determined by ITC and NMR assays are reported for all compounds featured in the challenge. All quantities are reported as point estimates ± statistical error obtained by error propagation. For *K*_*a*_ and Δ*H*, the reported uncertainties incorporate both the uncertainty in the ITC enthalpogram least-squares fit and an assumed 3% uncertainty in titrant concentration. A minimum least-squares fit uncertainty of 1% was assumed for fit errors reported by instrumentation as < 1%. Δ*G* and *T*Δ*S* and their uncertainties were obtained from the first two quantities. Some of the compounds in the CB8 guest set can be bound by their hosts with stoichiometries different than 1:1. For CB8-G1 and CB8-G4, which can form 1:2 (two hosts bound to the same guest) and 1:3 complexes with CB8, respectively, we report the thermodynamic quantities of only one of the equivalent binding events—the value used to calculate the statistics for challenge entries. For CB8-G12, we report the measurements of both the 1:1 (CB8-G12a) and the 2:1 (CB8-G12b) bound complexes. The original data can be found at https://github.com/MobleyLab/SAMPL6/tree/master/host_guest/Analysis/ExperimentalMeasurements/experimental_measurements.csv. Eventual updates or corrections to the data will be made available at the same URL, and anyone wishing to reuse the data should refer there. ^(*a*)^ Point estimate and uncertainties computed from the *K*_*a*_ measurements by error propagation. ^(*b*)^ All experiments were performed at 298 K. ^(*c*)^ The thermodynamic quantities given here represent the binding free energy and enthalpy of one of the 1/*n* equivalent binding events. ^(*d*)^ Units of M^‒2^.

| ID | $K_a$ ($M^{-1}$) | $\Delta G$ (kcal/mol) <sup>(a)</sup> | $\Delta H$ (kcal/mol) | $T\Delta S$ (kcal/mol) <sup>(b)</sup> | $n$ |
| --- | --- | --- | --- | --- | --- |
| OA-G0 | $(147 \pm 7) \times 10^2$ | $-5.68 \pm 0.03$ | $-4.8 \pm 0.2$ | $0.8 \pm 0.2$ | 1 |
| OA-G1 | $(26 \pm 1) \times 10^2$ | $-4.65 \pm 0.02$ | $-5.5 \pm 0.2$ | $-0.9 \pm 0.2$ | 1 |
| OA-G2 | $(140 \pm 6) \times 10^4$ | $-8.38 \pm 0.02$ | $-12.1 \pm 0.5$ | $-3.7 \pm 0.5$ | 1 |
| OA-G3 | $(62 \pm 2) \times 10^2$ | $-5.18 \pm 0.02$ | $-7.5 \pm 0.3$ | $-2.4 \pm 0.3$ | 1 |
| OA-G4 | $(164 \pm 7) \times 10^3$ | $-7.11 \pm 0.02$ | $-6.9 \pm 0.3$ | $0.2 \pm 0.3$ | 1 |
| OA-G5 | $(233 \pm 9) \times 10$ | $-4.59 \pm 0.02$ | $-5.3 \pm 0.2$ | $-0.7 \pm 0.2$ | 1 |
| OA-G6 | $(44 \pm 2) \times 10^2$ | $-4.97 \pm 0.02$ | $-5.3 \pm 0.2$ | $-0.3 \pm 0.2$ | 1 |
| OA-G7 | $(36 \pm 1) \times 10^3$ | $-6.22 \pm 0.02$ | $-7.4 \pm 0.3$ | $-1.2 \pm 0.3$ | 1 |
| TEMOA-G0 | $(28 \pm 1) \times 10^3$ | $-6.06 \pm 0.02$ | $-7.8 \pm 0.4$ | $-1.8 \pm 0.4$ | 1 |
| TEMOA-G1 | $(24 \pm 2) \times 10^3$ | $-5.97 \pm 0.04$ | $-8.2 \pm 0.6$ | $-2.3 \pm 0.6$ | 1 |
| TEMOA-G2 | $(98 \pm 4) \times 10^3$ | $-6.81 \pm 0.02$ | $-9.3 \pm 0.4$ | $-2.5 \pm 0.4$ | 1 |
| TEMOA-G3 | $(128 \pm 9) \times 10^2$ | $-5.60 \pm 0.04$ | $-8.9 \pm 0.4$ | $-3.2 \pm 0.4$ | 1 |
| TEMOA-G4 | $(51 \pm 2) \times 10^4$ | $-7.79 \pm 0.02$ | $-8.9 \pm 0.4$ | $-1.1 \pm 0.4$ | 1 |
| TEMOA-G5 | $(113 \pm 5) \times 10$ | $-4.16 \pm 0.02$ | $-8.0 \pm 0.3$ | $-3.8 \pm 0.3$ | 1 |
| TEMOA-G6 | $(91 \pm 5) \times 10^2$ | $-5.40 \pm 0.03$ | $-6.2 \pm 0.2$ | $-0.8 \pm 0.2$ | 1 |
| TEMOA-G7 | $(107 \pm 4) \times 10$ | $-4.13 \pm 0.02$ | $-8.3 \pm 0.3$ | $-4.2 \pm 0.3$ | 1 |
| CB8-G0 | $(81 \pm 6) \times 10^3$ | $-6.69 \pm 0.05$ | $-4.2 \pm 0.2$ | $2.5 \pm 0.2$ | 1 |
| CB8-G1<sup>(c)</sup> | $(40 \pm 3) \times 10^4$ | $-7.65 \pm 0.04$ | $-5.0 \pm 0.2$ | $2.6 \pm 0.2$ | 0.5 |
| CB8-G2 | $(41 \pm 4) \times 10^4$ | $-7.66 \pm 0.05$ | $-6.5 \pm 0.3$ | $1.2 \pm 0.3$ | 1 |
| CB8-G3 | $(53 \pm 5) \times 10^3$ | $-6.45 \pm 0.06$ | $-2.5 \pm 0.1$ | $4.0 \pm 0.2$ | 1 |
| CB8-G4<sup>(c)</sup> | $(51 \pm 4) \times 10^4$ | $-7.80 \pm 0.04$ | $-9.8 \pm 0.4$ | $-2.0 \pm 0.4$ | 0.33 |
| CB8-G5 | $(99 \pm 9) \times 10^4$ | $-8.18 \pm 0.05$ | $-3.2 \pm 0.1$ | $5.0 \pm 0.1$ | 1 |
| CB8-G6 | $(13 \pm 1) \times 10^5$ | $-8.34 \pm 0.05$ | $-5.7 \pm 0.2$ | $2.6 \pm 0.2$ | 1 |
| CB8-G7 | $(21 \pm 4) \times 10^6$ | $-10.0 \pm 0.1$ | $-6.5 \pm 0.3$ | $3.5 \pm 0.3$ | 1 |
| CB8-G8 | $(83 \pm 6) \times 10^8$ | $-13.50 \pm 0.04$ | $-14.4 \pm 0.6$ | $-0.9 \pm 0.6$ | 1 |
| CB8-G9 | $(23 \pm 3) \times 10^5$ | $-8.68 \pm 0.08$ | $-4.6 \pm 0.2$ | $4.0 \pm 0.2$ | 1 |
| CB8-G10 | $(10 \pm 1) \times 10^5$ | $-8.22 \pm 0.07$ | $-2.00 \pm 0.08$ | $6.2 \pm 0.1$ | 1 |
| CB8-G11 | $(50 \pm 4) \times 10^4$ | $-7.77 \pm 0.05$ | $-2.11 \pm 0.08$ | $5.7 \pm 0.1$ | 1 |
| CB8-G12a | $(167 \pm 9) \times 10^5$ | $-9.86 \pm 0.03$ | $-9.2 \pm 0.4$ | $0.7 \pm 0.4$ | 1 |
| CB8-G12b | $(146 \pm 6) \times 10^3$ <sup>(d)</sup> | $-7.05 \pm 0.02$ | $-4.8 \pm 0.2$ | $2.2 \pm 0.2$ | 2 |
| CB8-G13 | $(161 \pm 8) \times 10^3$ | $-7.11 \pm 0.03$ | $-6.8 \pm 0.3$ | $0.3 \pm 0.3$ | 1 |

**Figure 2.**
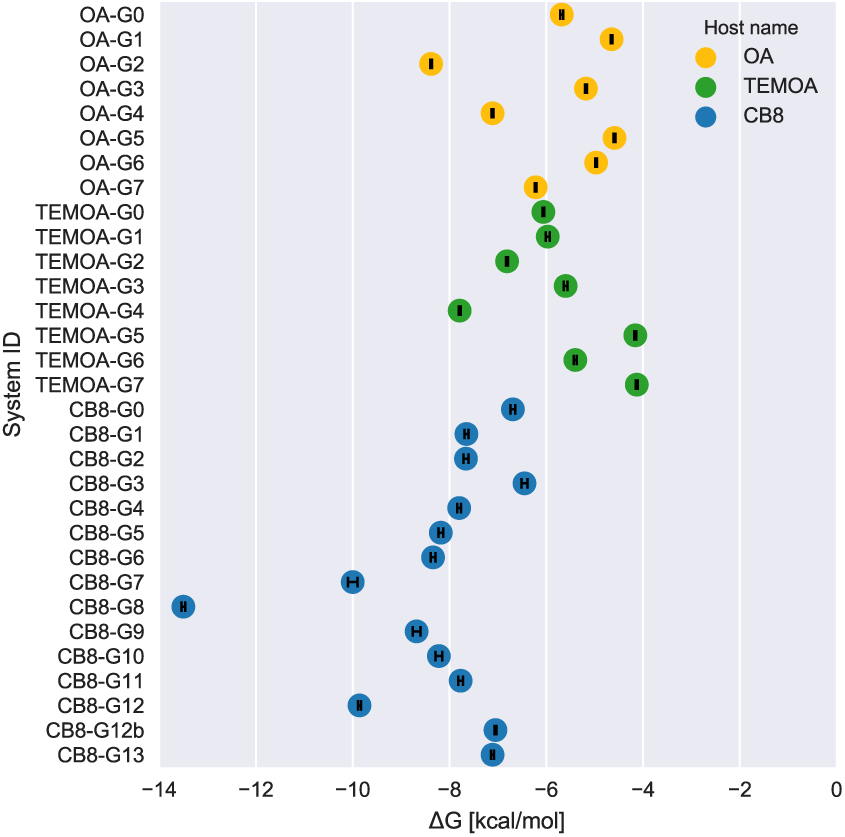
Overview of experimental binding affinities for all host-guest complexes in the SAMPL6 challenge set. Binding free energies (Δ*G*) measured via isothermal titration calorimetry (ITC) are shown (filled circles), along with experimental uncertainties denoting standard error of the mean (black error bars), for OA (yellow), TEMOA (green), and CB8 (blue) complexes.

## Methods

### Challenge design and logistics

#### Challenge timeline

On August 24th, 2017, we released in a publicly accessible GitHub repository [51] a brief description of the host-guest systems and the experimental methodology, together with the challenge directions, and input files in `mol2` and `sdf` formats for the three hosts and their guests. The instructions shared online included information about buffer concentrations, temperature, and pH used for the experiments. The participants were asked to submit their predicted absolute binding free energies and, optionally, binding enthalpies, along with a detailed description of the methodology and the software employed through the Drug Design Data Resource (D3R) website (https://drugdesigndata.org/about/sampl6) by January 19th, 2018. We also encouraged the inclusion of uncertainties and/or standard error of the mean (SEM) of the predictions when available. The results of the experimental assays were released on January 26th in the same GitHub repository. The challenge culminated in a conference held on February 22-23, 2018 in La Jolla, CA where the participants shared lessons learned from participating in the challenge after performing retrospective analysis of their data.

#### Bonus challenge

Three molecules in the CB8 guest sets, namely CB8-G11, CB8-G12, and CB8-G13, were proposed to participants as an optional bonus challenge since they were identified in advance to present some atypical difficulties for molecular modeling. In particular, the initial experimental data suggested both CB8-G11 and CB8-G12 to bind with 2:1 binding stoichiometry while CB8-G13 was deemed to be an especially challenging case for modeling due to the presence of a coordinated platinum atom, which is commonly not readily handled by classical force fields and usually requires larger basis sets for quantum mechanics (QM) calculations than those commonly employed with simple organic molecules. Further investigation after the start date of the challenge revealed an error in the calibration of a CB8 solution which affected the measurement of CB8-G11. After correcting the error, a 1:1 stoichiometry was recovered, and the experiment was repeated to validate the result. Unfortunately, the new data was obtained too late to send out a correction to all participants, so only six entries included predictions for this guest.

#### Preparation of standard input files

Standard input files for the three hosts were generated for the previous rounds of the SAMPL host-guest binding challenge and uploaded to the repository unchanged, while the guests’ atomic coordinates were generated from their SMILES string representation through the OMEGA library [46] in the OpenEye Toolkit (version 2017.Oct.b5) except for oxaliplatin (CB8-G13), which was generated with OpenBabel to handle the platinum atom. The compounds were then docked into their hosts with OpenEye’s FRED docking facility [72, 73]. Stereochemistry of the 3D structures recapitulated the stereochemistry of compounds assayed experimentally; experimental assays for chiral compounds were enantiopure except OA-G5, which was measured as a racemic mixture. For this molecule, we picked at random one of the two enantiomers under the assumption that the guest chirality (for this guest with a single chiral center) would not affect the binding free energy to an achiral host such as OA and TEMOA since the system otherwise contains no chiral centers. This information was included in the instructions when the challenge was released. Guest `mol2` files also included AM1-BCC point charges generated with the AM1-BCC charge engine in the Quacpac tool from the OpenEye toolkit [54, 55]. *Figure 1* shows the protonation state of the molecules as provided in the input files, which reflects the most likely protonation state as predicted by Epik [41, 102] from the Schrödinger Suite 2017-2 (Schrödinger) at experimental buffer pH (11.7 for OA and 7.4 for CB8). This resulted in all molecules possessing a net charge, with the exception of oxaliplatin and the CB8 host, which have no acidic or basic groups. Specifically, the eight carboxyl groups of OA and TEMOA were modeled as deprotonated and charged. The instructions stated clearly that the protonation and tautomeric states provided were not guaranteed to be optimal. In particular, participants in the bonus challenge were advised to treat CB8-G12 with care as, in its protonated state, the nitrogen proton could be placed so that the substituent was axial or equatorial. The latter solution was arbitrarily adopted by the tools used to generate the input files for CB8-G12.

### Statistical analysis of challenge entries

#### Performance statistics

We computed root mean square error (RMSE), mean signed error (ME), coefficient of determination (R^2^), and Kendall rank correlation coefficient (*τ*) comparing experimentally determined binding free energies with blinded participant free energy predictions.

The mean signed error (ME), which quantifies the bias in predictions, was computed as

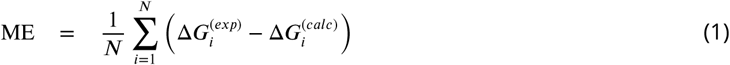

where 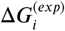 and 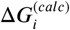 are the experimental measurement of the binding free energy and its computational prediction respectively for the *i*-th molecule, and *N* is the total number of molecules in the dataset. A positive ME reflects an overestimated binding free energy Δ*G* (or underestimated affinity *K*_*d*_ = *e*^‒*β*Δ*G*^ × (1 M).

Some of the methods appearing in SAMPL6 were also used in previous rounds of the same challenge to predict relative binding free energies of similar host-guest systems. In order to comment on the performance of these methods over sequential challenges, for which statistics on absolute free energies are not readily available, we computed a separate set of statistics defined as *offset statistics*, as opposed to the *absolute statistics* def ned above, in the same way they were reported in previous challenge overview papers. These statistics are computed identically to absolute statistics but by substituting 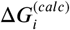 with

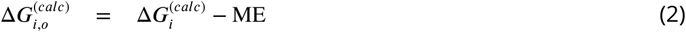

in the estimator expressions. The offset root mean square error computed from the 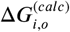 data points is termed RMSE_*o*_. It should be noted, however, that R^2^ and *τ* are invariant under a constant shift of the data points. For this reason, we will use the symbols R^2^ and *τ* both for the absolute and the offset correlation statistics.

Given the similarities of the two octa-acid hosts the set of their guest molecules, and that the large majority of the submitted methodologies were applied to both sets, we decided to report here the statistics computed using all the 16 predictions performed for OA and TEMOA (i.e., 8 predictions for each host). This merged set will be referred to as OA/TEMOA set in the rest of the work. The only method used to predict the binding free energies of the TEMOA set but not of the OA set was US-CGenFF (see *Table 2* for a schematic description of the methodology). We also decided to calculate separate statistics for the CB8 to highlight the general difference in performance between the predictions of the two host families. Statistics calculated on the two separate OA and TEMOA sets, as well as on the full dataset including CB8, OA, and TEMOA, are available on the GitHub repository (https://github.com/MobleyLab/SAMPL6/tree/master/host_guest/Analysis/Accuracy/).

**Table 2.** Summary of methodologies used by the participants in the SAMPL6 host-guest challenge. When a method uses multiple models (e.g., MM is used to generate the conformations to evaluate at the QM level in DFT(TPSS)-D3), only the energy and solvation models used for the final free energy prediction are listed. COSMO-RS: conductor-like screening model for real solvents [61]; DDM: double decoupling method [39]; FM: Force Matching [28]; FSDAM: Fast switching double annihilation method [92, 97] KMTISM: KECSA-Movable Type Implicit Solvation Model [131]; MD: molecular dynamics; MovTyp MovableType method [130]; PBSA: Poisson-Boltzmann surface area [106]; REST: replica exchange with solute torsional tempering [68, 71]; RFEC: relative free energy calculation; QM/MM: mixed quantum mechanics and molecular mechanics; SOMD: double annihilation or decoupling method performed with Sire/OpenMM6.3 software [27, Woods et al.]; SQM: semi-empirical quantum mechanics; US: umbrella sampling [119]; VSGB2.1: VSGB2.0 solvation model refitto OPLS2.1/3/3e [67]; ^(*a*)^ Alchemical calculations are flagged by (A). All of these are absolute free energy calculations except for the RFEC entries. ^(*b*)^ (E) and (I) denote explicit and implicit solvation models respectively. ^(*c*)^ The corrections based on previous experimental data either apply only an additive term (offset) or both an additive term and a multiplicative factor (linear). (*d*) Only a subset of the 25 movable type variations are included here. The four-letter suffix of each movable type submission is to be interpreted as following: first letter indicates the force field (G: GARF; K: KECSA), the second letter input structures (E: ensemble of structures from MD sampling; T: lowest energy structure during movable type scoring), the third letter is the number of states (1: only the complex is considered, 3: includes also the energy scores of host and guest in solution), and the fourth letter the type of experimental correction (L: linear; O: offset; N: no correction). ^(*e*)^ Both RFEC-GAFF2 and RFEC-QMMM report the results of relative free energy calculations. The offsets were determined from experimental data for similar OA or TEMOA guests. ^(*f*)^ SOMD submissions denoted with the *nobuffer* suffix include only the neutralizing counterions while the others add extra ions to model the buffer salt concentration. SOMD-A has no corrections. SOMD-B adds corrections for missing long-range dispersion interactions and for the flat-bottomed restraint to bring the ligand to standard state concentration. SOMD-D includes a linear correction fit to previously-available experimental data.

| Method ID<sup>(a)</sup> | Sampling | Energy model | Solvation model<sup>(b)</sup> | Experimental fit correction<sup>(c)</sup> | SAMPL6 reference |
| --- | --- | --- | --- | --- | --- |
| DDM-AMOEBA (A) | MD | AMOEBA | AMOEBA (E) | no |  |
| DDM-FM (A) | HREX; MD | Force-Matching/RESP | TIP3P (E) | no |  |
| DDM-FM-QMMM (A) | HREX; MD | Force-Matching/RESP;<br>DFT(B3LYP) | TIP3P (E) | no |  |
| DDM-GAFF (A) | MD | GAFF/AM1-BCC | TIP3P (E) | no |  |
| DFT(B3PW91) | MD; clustering | DFT(B3PW91) | SMD (I) | no |  |
| DFT(B3PW91)-D3 | MD; clustering | DFT(B3PW91)-D3 | SMD (I) | no |  |
| DFT(TPSS)-D3 | MD | DFT(TPSS)-D3 | COSMO-RS (I) | no |  |
| FSDAM (A) | REST; MD | GAFF2/AM1-BCC | TIP3P (E) | no |  |
| NULL | docking | OPLS3 | VSGB2.1 (I) | no |  |
| MMPBSA-GAFF | MD; clustering | GAFF/RESP | PBSA (I) | no |  |
| MovTyp-GE3N<sup>(d)</sup> | MD; clustering | GARF | KMTISM (I) | no |  |
| MovTyp-GE3O | MD; clustering | GARF | KMTISM (I) | offset |  |
| MovTyp-GE3L | MD; clustering | GARF | KMTISM (I) | linear |  |
| MovTyp-GT1N | MD; clustering | GARF | KMTISM (I) | no |  |
| MovTyp-GT1L | MD; clustering | GARF | KMTISM (I) | linear |  |
| MovTyp-KT1N | MD; clustering | KECSA | KMTISM (I) | no |  |
| MovTyp-KT1L | MD; clustering | KECSA | KMTISM (I) | linear |  |
| RFEC-GAFF2 (A)<sup>(e)</sup> | MD | GAFF2/RESP | TIP3P (E) | offset |  |
| RFEC-QMMM (A) | MD | GAFF2/RESP; PM6-DH+ | TIP3P (E) | offset |  |
| SQM(PM6-DH+) | MD | PM6-DH+ | COSMO-RS (I) | no |  |
| SOMD-A (A)<sup>(f)</sup> | MD | GAFF/AM1-BCC | TIP3P (E) | no |  |
| SOMD-A-nobuffer (A) | MD | GAFF/AM1-BCC | TIP3P (E) | no |  |
| SOMD-C (A) | MD | GAFF/AM1-BCC | TIP3P (E) | no |  |
| SOMD-C-nobuffer (A) | MD | GAFF/AM1-BCC | TIP3P (E) | no |  |
| SOMD-D (A) | MD | GAFF/AM1-BCC | TIP3P (E) | linear |  |
| SOMD-D-nobuffer (A) | MD | GAFF/AM1-BCC | TIP3P (E) | linear |  |
| US-CGenFF | MD | CGenFF | TIP3P (E) | no |  |
| US-GAFF | MD | GAFF/AM1-BCC | TIP3P (E) | no |  |
| US-GAFF-C | MD | GAFF/AM1-BCC | TIP3P (E) | linear |  |

Source: https://github.com/MobleyLab/SAMPL6/tree/master/host_guest/Analysis/Submissions

We generated bootstrap distributions of the statistics and computed 95-percentile bootstrap confidence intervals of the point estimates by generating 100 000 bootstrap samples through random sampling of the set of host-guest pairs with replacement. When the submission included SEMs for each prediction, we accounted for the statistical uncertainty in predictions by adding, for each bootstrap replicate, an additional Gaussian perturbation to the prediction with a standard deviation indicated by the SEM for that prediction.

#### Null model

In order to compare the results obtained by the participants to a simple model that can be evaluated with minimal effort, we computed the binding free energy predicted by MM-GBSA rescoring [40] using Prime [52, 53] with the OPLS3 forcefield [45] in the Schrödinger Suite 2018-1 (Schrödinger). We used the same docked poses provided in the input files that were shared with all the participants as the initial coordinates for all the calculations. All docked positions were minimized before being rescored with the OPLS3 force field and the VSGB2.1 solvent model. The only exception to this was CB8-G4, which was manually re-docked into the host, as the initial structure contained steric clashes that could not be relaxed by minimization, causing the predicted binding free energy to spike to an unreasonable value of +2443 kcal/mol.

## Results

We received 42 submissions for the OA guest set, 43 for TEMOA, and 34 for CB8, for a total of 119 submissions, from 10 different participants, 5 of whom uploaded predictions for the three compounds in the bonus challenge as well. Only two groups submitted enthalpy predictions, which makes it impractical to draw general conclusions about the state of the field regarding the reliability of enthalpy predictions. Moreover, the predictive performance was generally poor (see Supplementary *Figure 9*). The results of the enthalpy calculations are thus not discussed in detail here, but they are nevertheless available on the GitHub repository.

### Overview of the methodologies

Including the null model, 41 different methodologies were applied to one or more of the three datasets. In particular, the submissions included a total of 25 different variations of the movable type method exploring the effect of the input structures, the force field, the presence of conformational changes upon binding, and the introduction of previous experimental information on the free energy estimates. In order to facilitate the comparison among methods, we focus in this analysis on a representative subset of 7 different variations of the methodology. Supplementary *Figure 7* and Supplementary *Figure 8* show statistic bootstrap distributions and correlation plots for all the movable type free energy calculations submitted. As many of the methodologies are reported in detail elsewhere, in this section, we give a brief overview of the different strategies employed for the challenge to model the host-guest systems and estimate the binding free energies, and we leave the detailed descriptions of the various methodologies to the articles referenced in *Table 2*.

#### Modeling

The majority of the participants either used the docked poses provided in the input files or ran a separate docking program to generate the initial complex conformation for the calculations. In few cases, the starting configuration was found by manually placing the guest inside the host. Surprisingly, the most common solvent model used in classical simulations was still TIP3P [57], a water model parameterized by Jorgensen 35 years ago for use with a fixed-cutoff Monte Carlo code neglecting long-range dispersion interactions and omitting long-range electrostatics. The only other explicit water models used in this round of the challenge were the significantly more modern AMOEBA [96] and TIP4P-Ew [49] water models, which was used to sample conformations to evaluate at the QM level. Implicit solvent models were adopted only in MMPBSA and for the movable type and QM calculations. We observed more variability in the treatment of buffer salt concentrations despite the known importance of this element in affecting the binding predictions, which may reflect a lack of standard practices in the field. Some entries modeled the buffer ionic strength explicitly with Na+ and Cl-ions while others included only the neutralizing counterions or used a uniform neutralizing charge. One of the participating groups submitted multiple variants of the SOMD method either utilizing only neutralizing counterions or including additional ions simulating the ionic strength at experimental conditions, which makes it possible to directly assess the effect of this modeling decision on the selected host-guest systems.

Most methods employing classical force fields used GAFF [121 ] or GAFF2 (still under active development) with AM1-BCC [54, 55] or RESP [12] charges, which were usually derived at the Hartree-Fock or MP2 level of theory. Other approaches made use of the AMOEBA polarizable model [96], CGenFF [28] or force matching [120] starting from CGenFF parameters. The movable type calculations utilized either the KECSA [129] scoring algorithm or the more recently developed GARF [11]. Several submissions employed QM potentials at the semi-empirical PM6-DH+ [64, 100] or DFT level of theory either modeling the full host-guest system or in hybrid QM/MM approaches that treated quantum mechanically the guest only. DFT calculations employed B3LYP [13], B3PW91 [13], or TPSS [116] functionals and often the DFT-D3 dispersion correction [42].

#### Sampling and free energy prediction

All the challenge entries used MD to sample host-guest conformations; uses of docking were limited to preparation of initial bound geometries for subsequent simulations. This was also the case also for QM and movable type calculations, where samples generated from MD were in some cases clustered prior to quantum chemical energy evaluations. In a few cases, enhanced sampling techniques were used; in particular, the entries identified by DDM-FM and DDM-FM-QMM used Hamiltonian Replica Exchange (HREX) [113] as part of their double decoupling method (DDM) calculation [39] while Replica Exchange with Solute torsional Tempering (REST) [68, 71] was employed in FSDAM to generate from equilibrium the starting configurations for the fast switching protocol. Many groups used the double decoupling or the double annihilation method with purely classical force fields or with hybrid QM/MM potentials and either Bennett acceptance ratio (BAR) [15, 103] or the multistate Bennett acceptance ratio (MBAR) [104] to estimate free energies for the aggregated simulation data. Other classes of methodologies applied to this dataset include umbrella sampling (US) [119], movable type [130], MMPBSA [110], and free energy predictions based on QM calculations.

The repeat appearance of hosts chosen from the octa-acid and cucurbituril families as test systems for the SAMPL binding challenge, which reflects the continuous contribution of experimental data from the Gibb and Isaacs laboratories, led some groups to take advantage of previously available experimental data to improve their computational predictions. Several entries (e.g., SOMD-D, US-GAFF-C, and MovTyp-GE3L) were submitted with a linear^1^ correction of the form

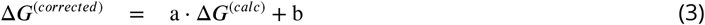

where the slope and offset coefficients (i.e., a and b respectively) were trained on data generated for previous rounds of the challenge. In some of the movable type calculations (e.g., MovTyp-GE3O), the coefficient a was fixed to unity and the training data used to determine a purely additive bias correction. Relatedly, RFEC-GAFF2 and RFEC-QMMM, which included predictions for the OA and TEMOA guest sets, calculated the relative binding free energy between the compound and determined the offsets necessary to obtain absolute free energy using binding measurements of similar OA and TEMOA guests.

### Submission performance statistics

As mentioned above, we present here the statistics obtained by the challenge entries on the CB8 dataset and the merged OA and TEMOA dataset with the exception of US-CGenFF, for which we received a submission for the TEMOA set only. Moreover, since only a minority of entries had predictions for the bonus challenge, we excluded CB8-G11, CB8-G12, and CB8-G13 when computing the statistics of all the methodologies in order to compare them on the same set of compounds. *Table 3* reports such statistics with 95-percentile confidence intervals and *Figure 4* show the statistics bootstrap distributions. Some of the methods were used to estimate the binding free energy of only one between the OA/TEMOA and the CB8 sets, and, as a consequence, some of the table entries are missing. For the methodologies that made predictions of the bonus compounds, we report the statistics obtained including them separately in *Table 4*. While it is difficult to isolate methods and models that performed very well across datasets and statistics, a few patterns emerged from comparing the different entries.

**Table 3.** Method performance statistics and bootstrap confidence intervals on OA/TEMOA and CB8 datasets. Root mean square error (RMSE), mean signed error (ME), coefficient of determination (R^2^), and Kendall correlation coefficient (τ) obtained by each methodology on the merged OA/TEMOA and the CB8 datasets. The only exception is US-CGenFF whose OA/TEMOA statistics were computed using only the TEMOA set since no submission was received for OA. Table entries are left blank for those methods that were applied to only one of the guest sets. The predictions performed for the bonus challenge guests were excluded when computing the statistics for the CB8 dataset. Each statistic is reported with bootstrap distribution mean (between parentheses) and 95-percentile bootstrap confidence interval (square brackets) obtained through 100 000 cycles of resampling with replacement. The standard errors of the mean of the predictions reported in the submissions are included in the confidence intervals. The original data for the combined OA/TEMOA and CB8 datasets can be found respectively at https://github.com/MobleyLab/SAMPL6/tree/master/host_guest/Analysis/Accuracy/OA-TEMOA/StatisticsTables/statistics.csv and https://github.com/MobleyLab/SAMPL6/tree/master/host_guest/Analysis/Accuracy/CB8-NOBONUS/StatisticsTables/statistics.csv. Eventual updates or corrections to the data will be made available at the same URL, and anyone wishing to reuse the data should refer there.

| Method | OA/TEMOA dataset |  |  |  | CB8 dataset (no bonus challenge) |  |  |  |
| --- | --- | --- | --- | --- | --- | --- | --- | --- |
| | RMSE | ME | $R^2$ | $\tau$ | RMSE | ME | $R^2$ | $\tau$ |
| DDM-AMOEBA | 2.2 (2.1) [1.3, 2.9] | -0.8 (-0.8) [-1.9, 0.1] | 0.1 (0.2) [0.0, 0.5] | -0.2 (-0.2) [-0.6, 0.2] | 3.9 (3.8) [1.5, 5.7] | 2.3 (2.3) [0.5, 4.3] | 0.1 (0.3) [0.0, 0.8] | 0.1 (0.1) [-0.5, 0.6] |
| DDM-FM |  |  |  |  | 4.7 (4.6) [3.4, 6.0] | 2.7 (2.7) [0.2, 4.8] | 0.4 (0.4) [0.0, 0.9] | 0.5 (0.5) [0.1, 0.9] |
| DDM-FM-QMMM |  |  |  |  | 5.5 (5.5) [4.0, 7.3] | 2.3 (2.3) [-1.1, 5.0] | 0.4 (0.5) [0.1, 0.8] | 0.6 (0.6) [0.1, 0.9] |
| DDM-GAFF | 3.4 (3.3) [2.1, 4.6] | 1.0 (1.0) [-0.7, 2.5] | 0.1 (0.3) [0.0, 0.8] | 0.4 (0.4) [-0.1, 0.8] | 7.2 (7.2) [5.6, 8.5] | 6.4 (6.4) [4.4, 8.2] | 0.3 (0.3) [0.0, 0.7] | 0.3 (0.3) [-0.2, 0.8] |
| DFT(B3PW91) | 16.7 (15.9) [8.7, 25.2] | 5.4 (5.4) [-3.4, 11.1] | 0.3 (0.3) [0.0, 0.7] | -0.2 (-0.2) [-0.7, 0.2] | 17.7 (17.5) [11.9, 22.9] | -14.8 (-14.8) [-20.6, -9.1] | 0.0 (0.1) [0.0, 0.7] | 0.3 (0.3) [-0.3, 0.8] |
| DFT(B3PW91)-D3 | 37.7 (37.7) [34.8, 40.1] | 35.3 (35.3) [27.8, 39.9] | 0.1 (0.3) [0.0, 0.8] | 0.4 (0.4) [-0.1, 0.9] | 36.6 (36.4) [28.7, 44.2] | 33.9 (33.9) [25.7, 42.0] | 0.0 (0.2) [0.0, 0.8] | -0.3 (-0.3) [-0.7, 0.2] |
| DFT(TPSS)-D3 | 3.1 (3.0) [2.3, 3.7] | -1.6 (-1.6) [-2.8, -0.2] | 0.5 (0.5) [0.1, 0.8] | 0.3 (0.4) [-0.1, 0.7] |  |  |  |  |
| FSDAM | 2.5 (2.5) [1.5, 3.3] | 0.8 (0.8) [-0.5, 1.9] | 0.5 (0.5) [0.0, 0.9] | 0.5 (0.5) [0.0, 0.9] |  |  |  |  |
| MMPBSA-GAFF | 7.0 (7.0) [5.4, 8.5] | 6.4 (6.4) [4.9, 7.9] | 0.8 (0.8) [0.6, 0.9] | 0.7 (0.6) [0.4, 0.8] | 17.9 (17.8) [13.9, 21.5] | 16.7 (16.7) [13.0, 20.6] | 0.0 (0.3) [0.0, 0.8] | -0.4 (-0.4) [-0.9, 0.1] |
| MovTyp-GE3L | 2.3 (2.3) [1.5, 3.1] | 1.9 (1.9) [1.2, 2.6] | 0.3 (0.4) [0.0, 0.8] | 0.3 (0.3) [-0.2, 0.7] | 5.8 (5.8) [4.5, 7.0] | 5.4 (5.4) [4.1, 6.7] | 0.3 (0.4) [0.1, 0.8] | 0.5 (0.5) [0.1, 0.8] |
| MovTyp-GE3N | 1.8 (1.8) [1.0, 2.6] | 1.2 (1.2) [0.5, 1.9] | 0.3 (0.4) [0.0, 0.8] | 0.3 (0.3) [-0.2, 0.7] | 4.7 (4.7) [3.5, 5.8] | -4.2 (-4.2) [-5.4, -2.8] | 0.3 (0.4) [0.1, 0.8] | 0.5 (0.5) [0.1, 0.8] |
| MovTyp-GE3O | 1.3 (1.3) [0.8, 1.8] | 0.8 (0.8) [0.3, 1.3] | 0.3 (0.4) [0.0, 0.8] | 0.3 (0.3) [-0.1, 0.7] | 5.3 (5.3) [4.2, 6.3] | 5.0 (5.0) [3.9, 6.1] | 0.3 (0.4) [0.1, 0.8] | 0.5 (0.5) [0.1, 0.8] |
| MovTyp-GT1L | 3.3 (3.3) [2.8, 3.8] | 3.2 (3.2) [2.7, 3.6] | 0.5 (0.5) [0.1, 0.8] | 0.4 (0.4) [-0.1, 0.8] | 5.4 (5.4) [4.1, 6.6] | -5.0 (-5.0) [-6.3, -3.7] | 0.4 (0.4) [0.1, 0.7] | 0.5 (0.5) [0.1, 0.9] |
| MovTyp-GT1N | 4.4 (4.4) [3.9, 4.9] | 4.3 (4.3) [3.8, 4.8] | 0.5 (0.5) [0.1, 0.8] | 0.4 (0.4) [-0.1, 0.8] | 2.0 (2.0) [1.3, 2.6] | -0.4 (-0.4) [-1.5, 0.8] | 0.4 (0.4) [0.1, 0.7] | 0.5 (0.5) [0.1, 0.9] |
| MovTyp-KT1L | 1.0 (0.9) [0.7, 1.2] | -0.5 (-0.5) [-0.9, -0.0] | 0.6 (0.6) [0.2, 0.8] | 0.4 (0.4) [-0.1, 0.7] | 2.9 (2.8) [1.7, 3.8] | 1.6 (1.6) [0.2, 3.0] | 0.1 (0.1) [0.0, 0.5] | 0.1 (0.1) [-0.4, 0.6] |
| MovTyp-KT1N | 2.9 (2.9) [2.4, 3.3] | 2.7 (2.7) [2.3, 3.2] | 0.5 (0.5) [0.1, 0.8] | 0.3 (0.3) [-0.1, 0.7] | 4.8 (4.8) [3.9, 5.6] | 4.4 (4.4) [3.3, 5.4] | 0.1 (0.1) [0.0, 0.5] | 0.1 (0.1) [-0.4, 0.6] |
| NULL | 26.3 (26.2) [23.3, 29.2] | 25.6 (25.6) [22.8, 28.6] | 0.6 (0.6) [0.2, 0.8] | 0.5 (0.6) [0.2, 0.8] | 17.6 (17.6) [14.2, 21.2] | 14.9 (14.9) [8.7, 19.9] | 0.0 (0.1) [0.0, 0.5] | -0.1 (-0.1) [-0.6, 0.5] |
| RFEC-GAFF2 | 1.5 (1.5) [1.2, 1.8] | -1.2 (-1.2) [-1.6, -0.7] | 0.7 (0.7) [0.3, 0.9] | 0.6 (0.6) [0.2, 0.9] |  |  |  |  |
| RFEC-QMMM | 1.6 (1.6) [1.3, 2.0] | -1.0 (-1.0) [-1.6, -0.3] | 0.8 (0.8) [0.6, 0.9] | 0.8 (0.7) [0.5, 0.9] |  |  |  |  |
| SOMD-A | 5.7 (5.7) [4.7, 6.6] | 5.4 (5.4) [4.5, 6.3] | 0.8 (0.8) [0.6, 0.9] | 0.8 (0.7) [0.4, 0.9] | 5.1 (5.2) [3.8, 6.8] | 4.4 (4.4) [2.6, 6.1] | 0.1 (0.2) [0.0, 0.8] | 0.1 (0.1) [-0.6, 0.7] |
| SOMD-A-nobuffer | 4.9 (4.9) [3.9, 5.9] | 4.5 (4.5) [3.6, 5.5] | 0.8 (0.8) [0.6, 0.9] | 0.7 (0.7) [0.4, 0.9] | 7.9 (7.9) [6.1, 9.7] | 7.3 (7.3) [5.5, 9.2] | 0.1 (0.2) [0.0, 0.7] | 0.1 (0.0) [-0.6, 0.6] |
| SOMD-C | 3.7 (3.7) [2.7, 4.5] | 3.2 (3.2) [2.4, 4.1] | 0.8 (0.8) [0.6, 0.9] | 0.7 (0.7) [0.4, 0.9] | 3.8 (3.9) [2.7, 5.5] | 2.8 (2.8) [1.0, 4.6] | 0.1 (0.2) [0.0, 0.8] | 0.1 (0.1) [-0.6, 0.7] |
| SOMD-C-nobuffer | 3.0 (3.0) [2.1, 3.9] | 2.4 (2.4) [1.4, 3.3] | 0.8 (0.8) [0.5, 0.9] | 0.7 (0.7) [0.4, 0.9] | 6.6 (6.6) [4.9, 8.4] | 6.0 (6.0) [4.1, 7.8] | 0.1 (0.2) [0.0, 0.7] | 0.1 (0.1) [-0.6, 0.6] |
| SOMD-D | 1.8 (1.7) [1.1, 2.4] | 1.0 (1.0) [0.4, 1.8] | 0.8 (0.8) [0.6, 0.9] | 0.7 (0.7) [0.4, 0.9] | 2.6 (2.6) [1.4, 3.8] | -1.8 (-1.8) [-3.1, -0.7] | 0.1 (0.2) [0.0, 0.8] | 0.1 (0.1) [-0.6, 0.7] |
| SOMD-D-nobuffer | 1.6 (1.6) [1.0, 2.2] | 0.3 (0.3) [-0.5, 1.1] | 0.8 (0.8) [0.5, 0.9] | 0.7 (0.7) [0.4, 0.9] | 1.9 (1.9) [1.2, 2.7] | -0.2 (-0.2) [-1.4, 1.0] | 0.1 (0.2) [0.0, 0.7] | 0.1 (0.0) [-0.6, 0.6] |
| SQM(PM6-DH+) | 2.7 (2.6) [1.8, 3.4] | 1.1 (1.1) [-0.0, 2.3] | 0.1 (0.2) [0.0, 0.7] | 0.3 (0.3) [-0.2, 0.8] |  |  |  |  |
| US-CGenFF | 1.3 (1.4) [0.7, 2.1] | -0.1 (-0.1) [-1.1, 1.0] | 0.5 (0.5) [0.0, 1.0] | 0.4 (0.4) [-0.4, 1.0] |  |  |  |  |
| US-GAFF | 2.9 (2.9) [2.2, 3.5] | 2.5 (2.5) [1.8, 3.2] | 0.9 (0.8) [0.6, 1.0] | 0.7 (0.7) [0.4, 0.9] | 8.0 (7.9) [4.9, 11.0] | 6.7 (6.7) [4.3, 9.5] | 0.0 (0.2) [0.0, 0.6] | -0.1 (-0.1) [-0.6, 0.5] |
| US-GAFF-C | 1.0 (0.9) [0.6, 1.2] | -0.5 (-0.5) [-0.9, -0.1] | 0.9 (0.8) [0.6, 1.0] | 0.7 (0.7) [0.4, 0.9] | 3.5 (3.4) [1.4, 5.2] | 1.6 (1.6) [-0.1, 3.6] | 0.0 (0.2) [0.0, 0.6] | -0.1 (-0.1) [-0.6, 0.5] |

**Figure 3.**
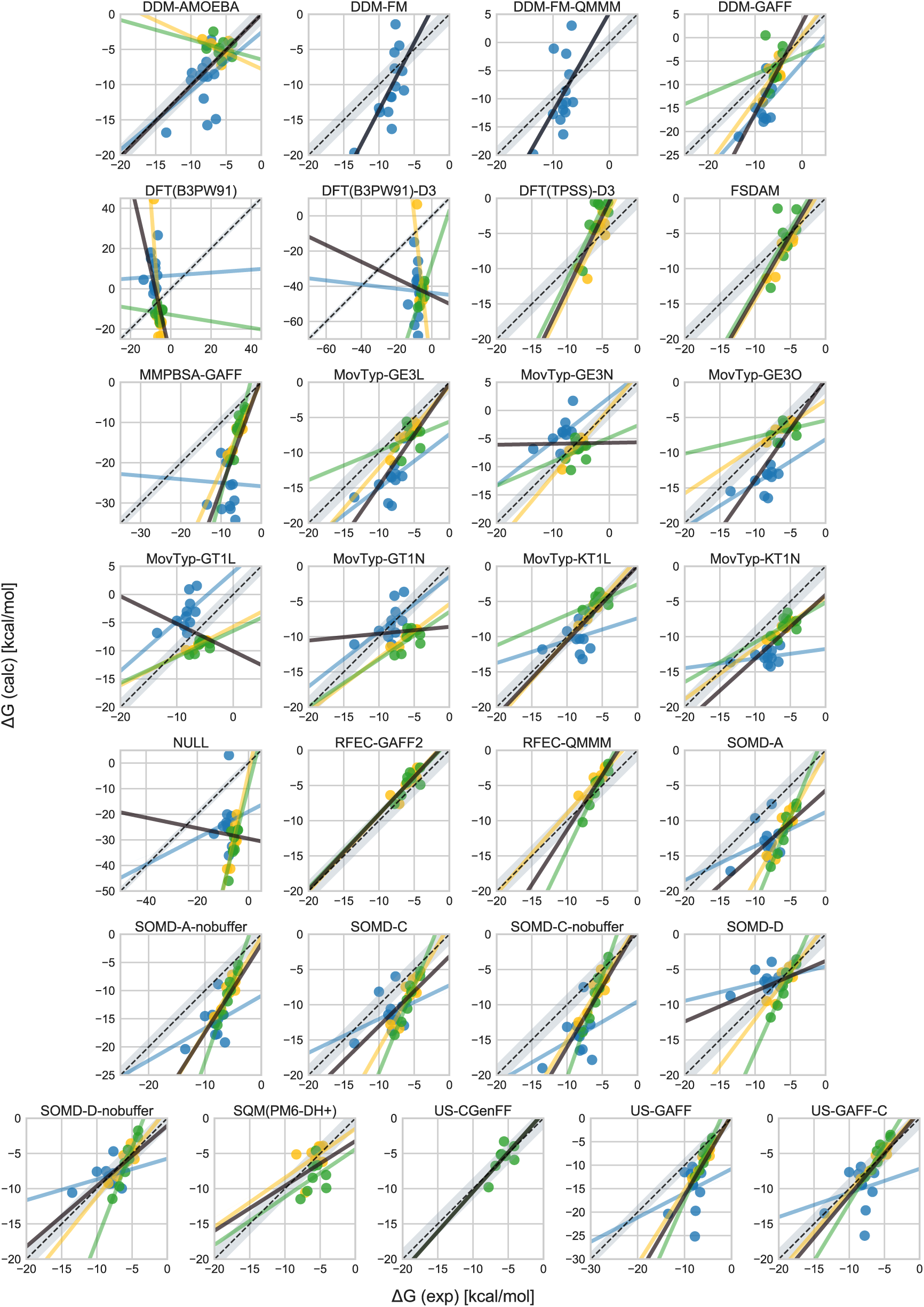
Free energy correlation plots obtained by the methods on the three host-guest sets. Scatter plots showing the experimental measurements of the host-guest binding free energies (horizontal axis) against the methods’ predictions on the OA (yellow), TEMOA (green), and CB8 (blue) guest sets with the respective regression lines of the same color. The solid black line is the regression line obtained by using all the data points. The gray shaded area represent the points within 1.5 kcal/mol from the diagonal (dashed black line). Only a representative subset of the movable type calculations results are shown. See Supplementary *Figure 7* for the free energy correlation plots of all the movable type predictions.

**Figure 4.**
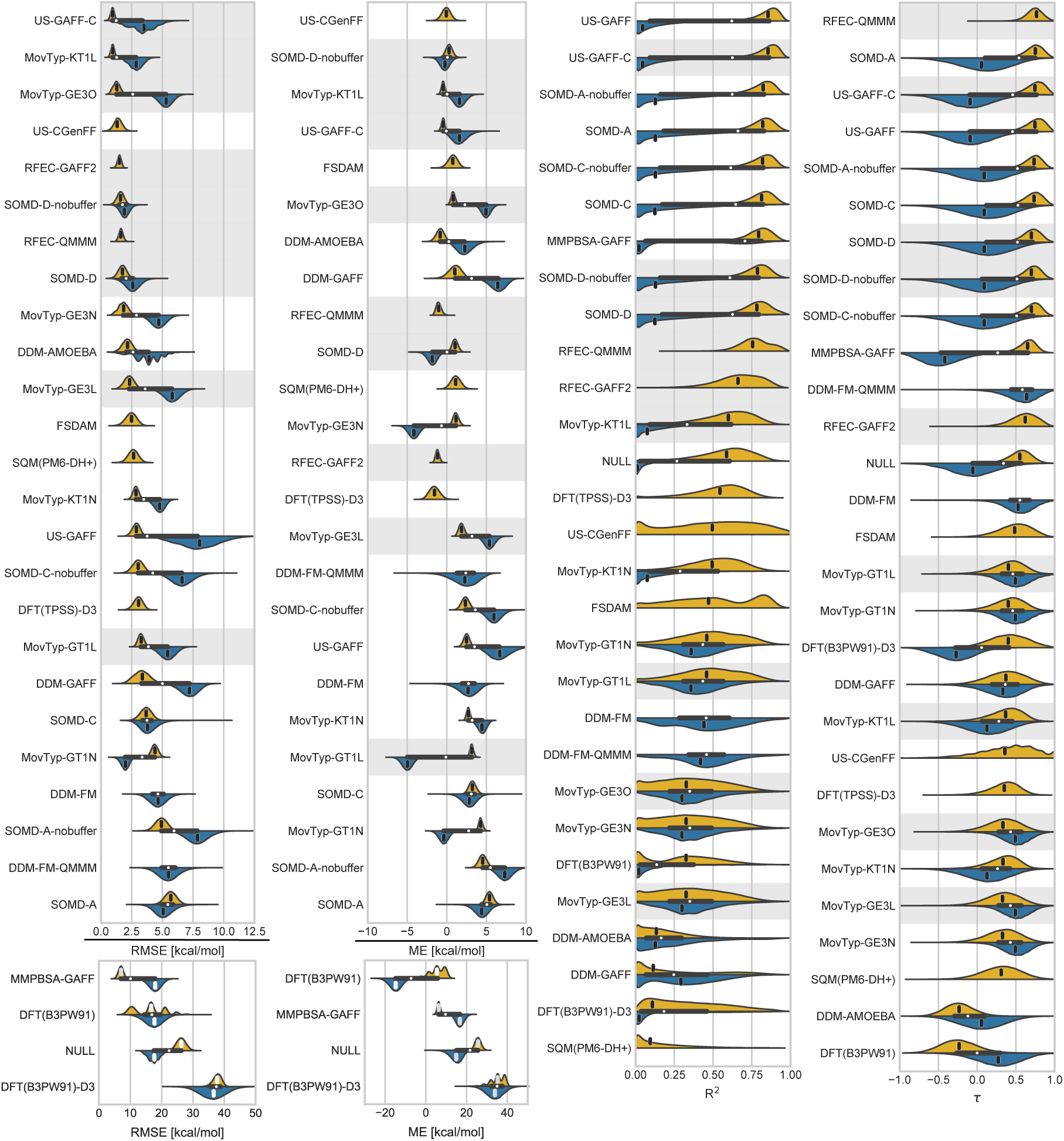
Bootstrap distribution of the methods performance statistics. Bootstrap distributions of root mean square error (RMSE), mean signed error (ME), coefficient of determination (R^2^) and Kendall rank correlation coefficient (*τ*). For each methodology and statistic, two distributions are shown for the merged OA/TEMOA set (yellow, pointing upwards) and the CB8 set excluding the bonus challenge compounds (blue, downwards). The black horizontal box between the two distributions of each method shows the median (white circle) and interquartile range (box extremes) of the overall distribution of statistics (i.e., pooling together the OA/TEMOA and CB8 statistic distributions). The short vertical segment in each distribution is the statistic computed using all the data. The distributions of the methods that incorporate previous experimental data into the computational prediction are highlighted in gray. Methodologies are ordered using the statistics computed on the OA/TEMOA set, unless only data for the CB8 set was submitted (e.g., DDM-FM), in which case the CB8 set statistic was used to determine the order. Only a representative subset of the movable type calculations results are shown. See Supplementary *Figure 8* for the bootstrap distributions including all the movable type submissions.

**Table 4.** Performance statistics including the bonus challenge molecules. Root mean square error (RMSE), mean signed error (ME), coefficient of determination (R^2^), and Kendall correlation coefficient (*τ*) obtained by all methods applied to the bonus challenge on the full CB8 set (left super column), including the three bonus molecules. Statistics computed excluding the bonus molecules are reported again here (right super column) for easy comparison. Bootstrap distribution mean and 95-percentile confidence intervals are reported between parentheses and square brackets respectively.

| Method | CB8 dataset (with bonus challenge) |  |  |  | CB8 dataset (no bonus challenge) |  |  |  |
| --- | --- | --- | --- | --- | --- | --- | --- | --- |
| | RMSE | ME | $R^2$ | $\tau$ | RMSE | ME | $R^2$ | $\tau$ |
| DDM-AMOEBA | 3.7 (3.6) [1.9, 5.2] | 1.2 (1.2) [-0.6, 3.1] | 0.1 (0.2) [0.0, 0.7] | 0.1 (0.1) [-0.3, 0.6] | 3.9 (3.8) [1.5, 5.7] | 2.3 (2.3) [0.5, 4.3] | 0.1 (0.3) [0.0, 0.8] | 0.1 (0.1) [-0.5, 0.6] |
| DDM-FM | 4.3 (4.4) [3.1, 5.6] | 2.2 (2.2) [0.0, 4.2] | 0.5 (0.5) [0.1, 0.8] | 0.5 (0.5) [0.2, 0.8] | 4.7 (4.6) [3.4, 6.0] | 2.7 (2.7) [0.2, 4.8] | 0.4 (0.4) [0.0, 0.9] | 0.5 (0.5) [0.1, 0.9] |
| DDM-FM-QMMM | 5.4 (5.5) [3.8, 7.3] | 1.1 (1.1) [-2.0, 3.8] | 0.3 (0.3) [0.0, 0.7] | 0.5 (0.4) [-0.0, 0.8] | 5.5 (5.5) [4.0, 7.3] | 2.3 (2.3) [-1.1, 5.0] | 0.4 (0.5) [0.1, 0.8] | 0.6 (0.6) [0.1, 0.9] |
| DFT(B3PW91) | 17.3 (17.1) [12.1, 21.8] | -14.4 (-14.4) [-19.5, -9.5] | 0.0 (0.1) [0.0, 0.5] | 0.1 (0.1) [-0.4, 0.5] | 17.7 (17.5) [11.9, 22.9] | -14.8 (-14.8) [-20.6, -9.1] | 0.0 (0.1) [0.0, 0.7] | 0.3 (0.3) [-0.3, 0.8] |
| DFT(B3PW91)-D3 | 37.0 (36.8) [29.9, 43.5] | 34.3 (34.3) [27.0, 41.4] | 0.0 (0.1) [0.0, 0.5] | -0.1 (-0.1) [-0.6, 0.2] | 36.6 (36.4) [28.7, 44.2] | 33.9 (33.9) [25.7, 42.0] | 0.0 (0.2) [0.0, 0.8] | -0.3 (-0.3) [-0.7, 0.2] |
| MMPBSA-GAFF | 17.8 (17.7) [14.5, 20.8] | 16.7 (16.7) [13.5, 19.9] | 0.0 (0.1) [0.0, 0.7] | -0.2 (-0.2) [-0.7, 0.2] | 17.9 (17.8) [13.9, 21.5] | 16.7 (16.7) [13.0, 20.6] | 0.0 (0.3) [0.0, 0.8] | -0.4 (-0.4) [-0.9, 0.1] |

Data Source: https://github.com/MobleyLab/SAMPL6/tree/master/host_guest/Analysis

#### Challenge entries generally performed better on OA/TEMOA than CB8

In general, the CB8 guest set proved to be more challenging than the OA/TEMOA set both in terms of error and correlation statistics. It is rarely the case that the same method scored better statistics on the former set, and only MovTyp-GT1N does so with statistical significance while the opposite can be observed relatively often. *Figure 5*-A shows the root mean square error (RMSE) and mean signed error (ME) with 95-percentile bootstrap confidence interval computed for each molecule using the ten methods that scored best in RMSE statistics in the merged OA/TEMOA set or the CB8 set (excluding the bonus challenge), which formed a set of 14 different techniques employing GAFF and GAFF2 [121], CGenFF [120], force matching [28], AMOEBA [96], and QM/MM potentials using DFT(B3LYP) [13] or PM6-DH+ [64, 100]. These top ten methods performed poorly on eight out of the eleven CB8 compounds, and while confidence intervals for all the statistics are generally large, they also performed significantly worse on several CB8 guests than the OA/TEMOA ligands they accurately predicted affinities for. This loss of accuracy seems to be fairly consistent across models and methodologies, but the data is not sufficient to determine the exact cause of this behavior (e.g., force field parameters, the generally larger dimensions of the CB8 guests, protonation states). However, the results of the related SAMPL6 SAMPLing challenge does suggest that properly accounting for slow conformational dynamics for some of the CB8 guests may require longer simulation times than for the OA compounds [51], which may have contributed to poorer performance over the OA set. Moreover, explicitly modeling the buffer salt concentration in SOMD significantly reduced the difference in error on the two guest sets (compare SOMD-C with SOMD-C-nobuffer), albeit without a commensurate improvement in correlation statistics, so the issue of missing chemical effects may also have role.

**Figure 5.**
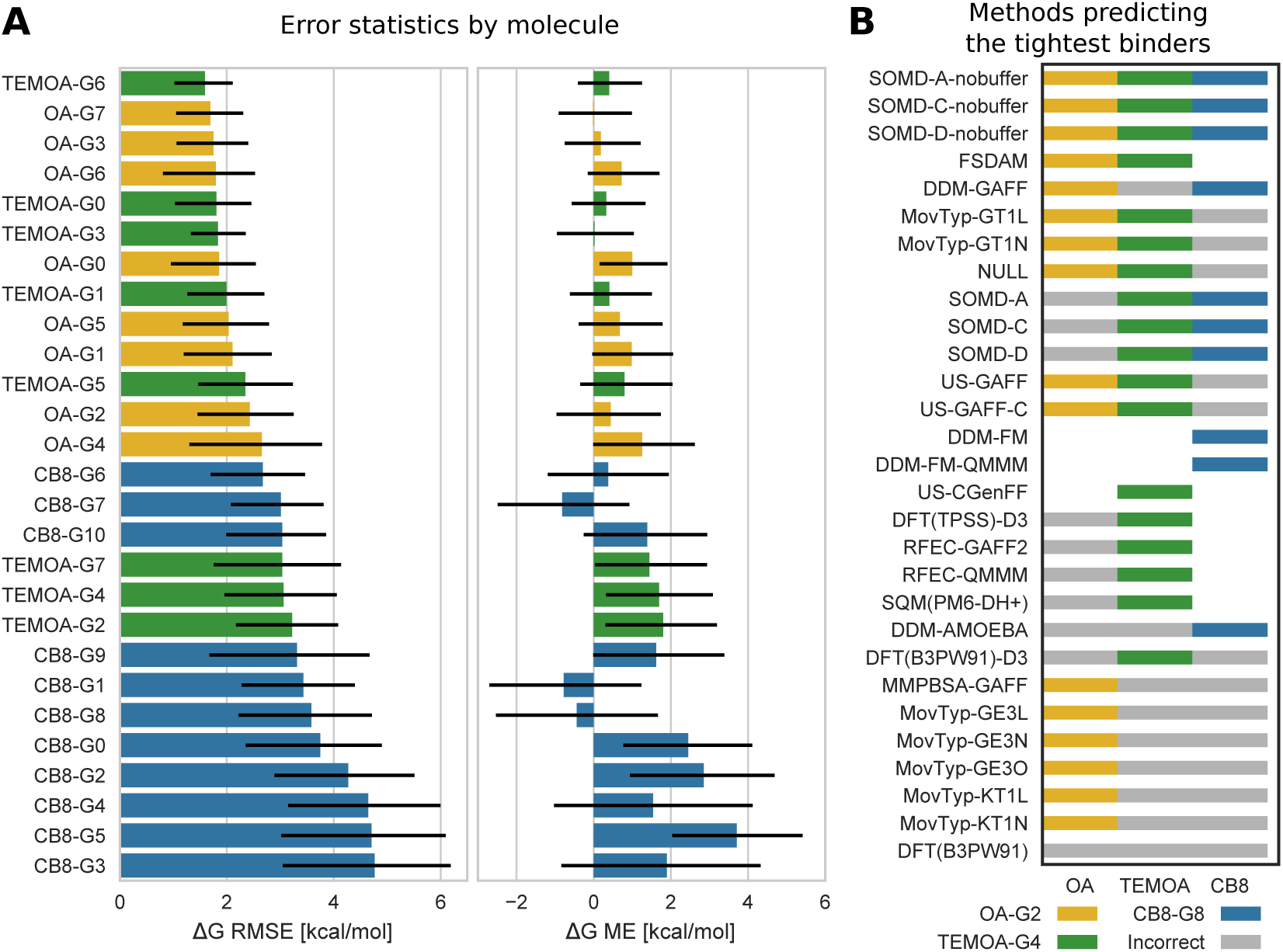
Free energy error statistics by molecule and tightest binders ranking. (**A**) Root mean square error (RMSE) and mean signed error (ME) computed using the ten methodologies with the lowest RMSE on the merged OA/TEMOA and CB8 datasets (excluding bonus challenge compounds) for all guests binding to OA (yellow), TEMOA (green), and CB8 (blue). Error bars represent 95-percentile bootstrap confidence intervals. (**B**) Ranking of the tightest binder of each host-guest dataset for all methods. Methods that correctly predicted OA-G2, TEMOA-G4, and CB8-G8 to be the tightest binders of the OA (yellow), TEMOA (green), and CB8 (blue) guest sets respectively are marked by a colored cell. A gray cell is shown when the method incorrectly predicted the tightest binder, and a white space is left if no submissions were received for that method and guest set. The methods are ordered by the number of correctly ranked tightest binders in the three guest sets.

The same trend appears when examining the performance of methods in correctly predicting the tightest binder of the three guest sets *Figure 5*-B. About 61% and 66% of the methods correctly ranked OA-G2 and TEMOA-G4 as the tightest complexes in their respective sets, while CB8-G8 was correctly classified in only about 43% of the cases. In particular, the latter observation is interesting when considering that the binding free energy of G8 to CB8 is 3.5 kcal/mol greater than the second tightest binder (G7), despite the structural similarities between both guests. It is also worth mentioning that SOMD method was the only methodology that correctly ranked the tightest binder of the three separate guest sets, although the prediction that G8 was the highest affinity guest for CB8 did not hold when buffer salt conditions were modeled explicitly.

#### Linear corrections fit to prior experimental data can reduce error without improving correlation

Nine of the entries represented in *Figure 4* incorporate fits to prior experimental data with the goal of either improving the computationally-predicted affinities or determining the offset necessary to convert relative free energy estimates into absolute binding affinities. It should be noted that a constant offset or multiplicative factor modifying all data points *cannot* alter the R^2^ statistic besides correcting an inverse correlation, and they can change τ only if the transformation is such that the ranking of at least two data points is switched, which a single linear transformation with positive slope cannot do. However, since some of the entries fit distinct correction terms for OA and TEMOA guests, correlation statistics for the combined OA/TEMOA set were affected (see, e.g., Supplementary *Figure 8* results for SOMD-C and SOMD-D, MovTyp-GE3N and MovTyp-GE3S). We can thus observe the effects of the linear transformations trained on experimental data on both the error and correlation statistics.

The corrections were generally successful in reducing RMSE. Among the top 10 methods scoring the lowest RMSE on the OA/TEMOA set, seven employ a correction. Moreover, when considering multiple submissions of the same technique that differ only in whether a fit to prior experimental data was included, the entry with the lowest RMSE incorporates experimental data in every case. However, the results are less consistent when considering the CB8 guest set. The trend is the same for the SOMD, US-GAFF, and MovTyp submissions that used the KECSA potential, but it is reversed for the majority of the MovTyp submissions employing the GARF energy model (see also Supplementary *Figure 8*). It should be noted that many of the MovTyp corrections were trained on a dataset that pooled binding measurements of OA, TEMOA, and CB8 guests, so it is possible that the approach failed to generalize when the methodology was affected by a systematic error of opposite sign on the OA/TEMOA and CB8 sets (see *Figure 3*). The methods that scored best (in terms of lowest RMSE) are US-GAFF-C for OA/TEMOA, and SOMD-D-nobuffer for CB8; excluding methods utilizing fits to experimental data, US-CGenFF and MovTyp-GT1N have the lowest RMSE on the OA/TEMOA and CB8 sets, respectively.

On the other hand, integrating prior experimental data did not appreciably impact correlation statistics, and the same methods with or without experimental correction show very similar R^2^ and *τ* bootstrap distributions. It is true that the initial performance of these methods without the experiment-based correction on the separated OA and TEMOA sets was relatively similar, thus leaving a small margin of improvement for this type of correction to reduce the data variance around the regression line and increasing R^2^. However, comparing the statistics computed pooling together the OA/TEMOA and CB8 predictions, which displayed very different correlation statistics, did not show any significant improvement (data not shown). In fact, R^2^ for the SOMD-C calculations decreased from 0.47 [0.09,0.78] to 0.18 [0.01,0.48] when incorporating the experimental correction in SOMD-D, despite the expected drop in RMSE, and a similar observation can be made for SOMD-D-nobuffer and the τ statistic.

#### GAFF/AM1-BCC and TIP3P consistently overestimated the host-guest binding affinities

Several entries used GAFF to parameterize the host-guest systems with AM1-BCC charges and TIP3P water molecules (i.e., SOMD, US-GAFF, DDM-GAFF) so it is possible to make relatively general observations about the performance of this model. Firstly, if we ignore the submissions that employ an experiment-based correction, every single method in this group predicted tighter binding than what supported by experiments with both the OA/TEMOA and the CB8 sets. This observation extends to MMPBSA-GAFF as well, which still used GAFF but with RESP charges and the implicit PBSA solvent model, but many of the methodologies that entered the challenge display a similar systematic error (see also ME in *Figure 5*), although GAFF is the only force field that was independently adopted by multiple groups and used with various classes of techniques.

Secondly, while error statistics vary substantially among GAFF entries, the correlation statistics are quite similar. Most of these are among the best-performing methods for the OA/TEMOA set, with *τ* ranging between 0.7–0.8, despite showing poor correlations on the CB8 set. The main exception to this pattern is given by DDM-GAFF, which shows moderate correlations for both datasets. The reason for this is not entirely clear, as the methodology adopted for DDM-GAFF entry is very similar to SOMD-C-nobuffer. Their main difference appears to lie in their treatment of long-range electrostatics, with SOMD using reaction field electrostatics [117] and DDM-GAFF using PME [29], as well as the use of restraints, with SOMD employing a single flat-bottom restraint to keep the guest in the host’s cavity and DDM-GAFF restraining the relative orientation of the guest by means of harmonic restraining potentials applied to one distance, two angles, and three torsions.

#### Models accounting for polarization did not perform significantly better than point charge models

Several of the entries adopted explicit model of electrostatic polarization through either QM potentials or the AMOEBA force field. Two groups submitted predictions obtained both a point-charge force field that were corrected with the free energy of moving to a QM/MM potential. This is the case of RFEC-QMMM and DDM-FM-QMMM, both of which included only the guest in the QM region using PM6-DH+ and DFT(B3LYP) respectively. In both cases, when compared to the pure MM model, the correlation slightly increased, although this difference was not statistically significant. Notably, RFEC-QMMM and DDM-FM-QMMM scored the top *τ* for the OA/TEMOA and CB8 set respectively.

On the other hand, calculations based on the polarizable AMOEBA force field or pure QM potentials were generally outperformed by point charge force-fields and QM/MM models in terms of correlation with experimental data. However, when limiting the comparison to methods that did not include a linear correction fitted on previous experimental data, SQM(PM6-DH+), DFT(TPSS)-D3, and in particular DDM-AMOEBA obtained a relatively low RMSE in spite of their poor correlation with experimental data. It is of interest to note that SQM(PM6-DH+) and DFT(TPSS)-D3 performed similarly. Indeed, the two methodologies were submitted by the same group and differ only by the potential function used to compute the energy of the complex on a set of configurations sampled with MD. SQM(PM6-DH+) scored a slightly lower RMSE and DFT(TPSS)-D3 obtained slightly higher correlation statistics, but the difference is not statistically significant in either case. The data, however, seems to suggest opposite tendencies of the two models in regard to the bias, with SQM(PM6-DH+) and DFT(TPSS)-D3 overestimating and underestimating the binding affinity of the OA/TEMOA guest set respectively. Similarly, DFT(B3PW91) and DFT(B3PW91)-D3 differ exclusively by the addition of the dispersion correction, which, surprisingly, significantly worsen the error for both guest sets.

#### Comparison to null model

The vast majority of the entries statistically outperformed the MMGBSA calculation we used as a null model. Surprisingly, while the null model correlation on the CB8 set was objectively poor (R^2^ = 0.0 [0.0, 0.5], *τ* = -0.1 [-0.6, 0.5]), the R^2^ and *τ* statistics obtained by the MMGBSA null model on the OA/TEMOA set was comparable to more expensive methods and, in fact, surpassed many of the challenge entries (*Table 3*). Nevertheless, the MMGBSA null model was in general poorly accurate in terms of RMSE. We note the difference of our null model with the MMPBSA-GAFF, which generally performed better than MMGBSA on the OA/TEMOA guest set but similarly or slightly worse on the CB8 set. Besides differences in solvent model (i.e., Generalized Born and Poisson-Boltzmann respectively), the former used OPLS3 to rescore a single docked pose, while the second one used GAFF and molecular dynamics to collect samples that were subsequently clustered for the purpose of rescoring.

### Bonus challenge

The platinum atom in CB8-G13 required particular attention during parameterization as this atom is not customarily handled by general small molecule force fields. Even in the case of DFT(B3PW91) and DFT(B3PW91)-D3, the configurations used for the QM calculations were generated by classical molecular dynamics requiring empirical parameters. In general, all the participants to the bonus challenge relied on DFT-level quantum mechanics calculation to address the problem. In MMPBSA-GAFF, DFT(B3PW91), and DFT(B3PW91)-D3, Mulliken charges were generated from DFT(B3LYP), which were subsequently used to determine AM1-BCC charges. A different approach was adopted in DDM-FM-QMMM in which the platinum was substituted by palladium, and the conformations necessary to the force matching parameterization procedure were obtained by MNDO(d) dynamics.

All groups participating to the bonus challenge submitted 1:1 complex predictions also for CB8-G11 and CB8-G12, for which the initial experimental data suggested the possibility of 2:1 complexes (two guests simultaneously bound to one host). This later turned out to be correct only for CB8-G12, and several groups reported to have computationally tested the hypothesis for CB8-G11 with the correct outcome. DDM-AMOEBA was used to estimate affinity of both the 1:1 and 2:1 complexes, but in the end the first one was used in the submission as the two predicted binding free energies differed by only 0.1 kcal/mol. Accordingly, we used the experimental measurement determined for the first binding event to compute the statistics (CB8-G12a in *Table 1*).

Summary statistics incorporating bonus challenge compounds are reported in *Table 4*. Although the RMSE generally improves in most cases, it should be noted that this effect varies greatly across the three molecules, and this improvement is mainly due to CB8-G11, whose predictions are regularly much closer to the experimental measurement than the estimates provided for the other two compounds.

### Comparison to previous rounds of the SAMPL host-guest binding challenge

Since previous rounds of the host-guest binding challenge featured identical or similar hosts to those tested in SAMPL6, it is possible to compare earlier results and observe the evolution of methodological performance.

#### Accuracy improvements over SAMPL5 for OA/TEMOA were driven by fits to prior experimental data

SAMPL5 featured a set of compounds binding to both OA and TEMOA, which will be referred in the following as the OA/TEMOA-5 set to differentiate it from the combined OA/TEMOA set used in this round of the challenge. In the top row of *Figure 6*-A, we show median and fitted distributions of the RMSE and R^2^ statistics taken from the SAMPL5 overview paper [127] together with the results from SAMPL6. OA was used as a test system in SAMPL4 as well, but in this case, only relative free energy predictions were submitted so we cannot draw a direct comparison. Prediction accuracy displays a slight improvement of the median RMSE from the previous round from 3.00 [2.70, 3.60] kcal/mol to 2.76 [1.85, 3.28] kcal/mol (95-percentile bootstrap confidence intervals of the medians not shown in *Figure 6*-A). However, this change seems to be entirely driven by the methods employing experiment-based fit corrections since removing them results in a median RMSE that is essentially identical to SAMPL5. The data raises the question of whether the field is hitting the accuracy limit of current general force fields.

**Figure 6.**
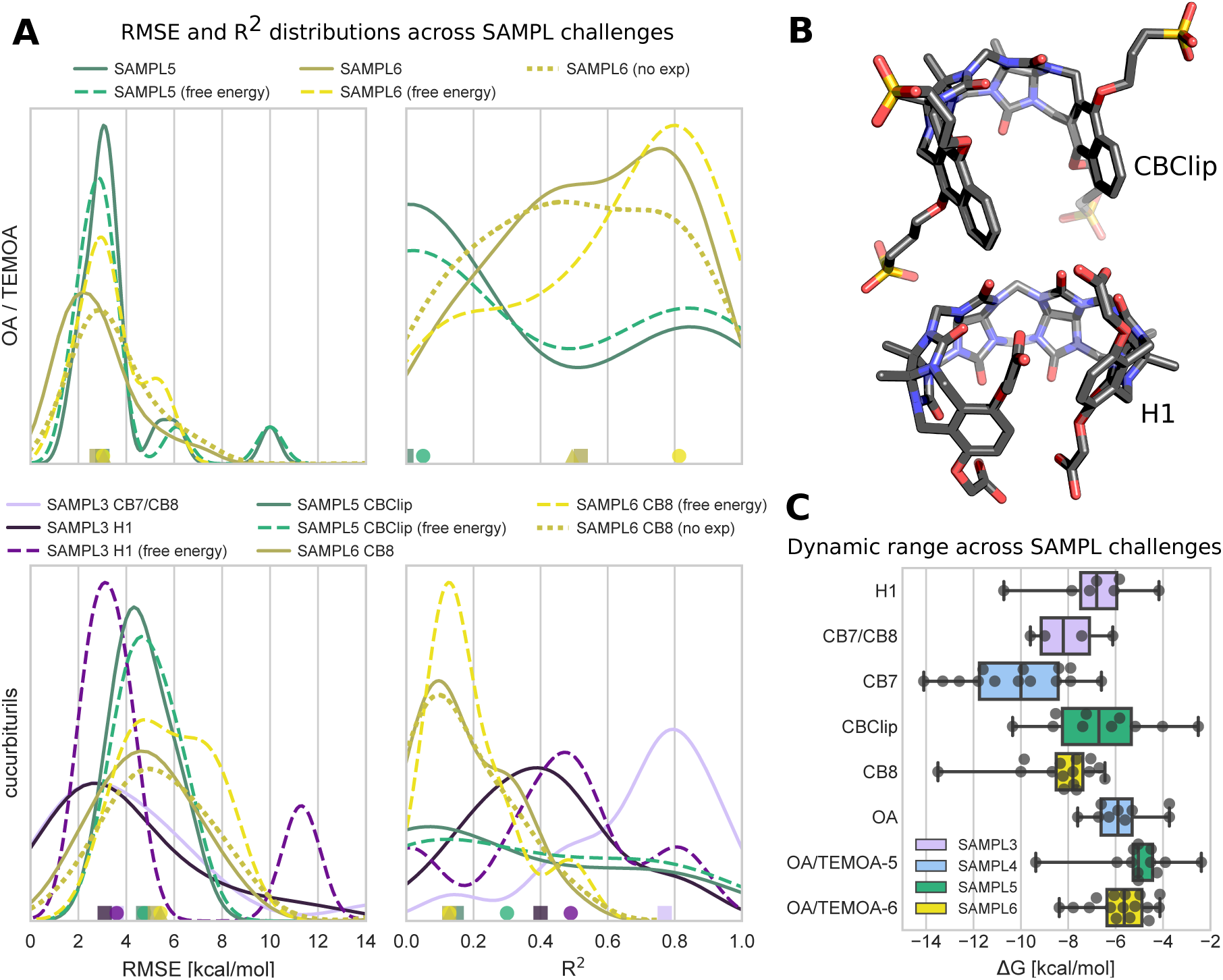
CB analogues and distribution of RMSE and R^2^ achieved by methods in SAMPL3 and SAMPL5. (**A**) Probability distribution fitting of root mean square error (RMSE, left column) and coefficient of determination (R^2^, right column) achieved by all the methods entering the SAMPL6 (yellow), SAMPL5 (green), and SAMPL3 (purple) challenge. Statistics for SAMPL4 are not shown in the panel because the subject of the challenge was confined to relative binding affinity predictions. The markers on the x-axis indicate the medians of the distributions. Distributions are shown for all the methods entering the challenge (solid line, square marker), excluding the SAMPL6 entries that used previous experimental data (dotted line, triangle marker), or isolating alchemical and potential of mean force methodologies that did not use an experiment-based correction (dashed line, circle marker). The RMSE axis is truncated to 14 kcal/mol, and a few outlier submissions are not shown. The data shows an essentially identical median RMSE and an increased median correlation on the combined OA/TEMOA guest sets (top row) with respect to the previous round of the challenge. The comparison of the results to different sets of guests binding few cucurbit[n]uril and cucurbit[n]uril-like hosts appearing in SAMPL3 and SAMPL5 (bottom row) shows instead a deteriorated performance in the most recent round of the challenge, which is likely explained by the major complexity of the SAMPL6 C8 guest set. (**B**) Three-dimensional structures in stick view of the CBClip (top) and H1 (bottom) hosts featuring in SAMPL5 and SAMPL3 respectively. Carbon atoms are represented in gray, nitrogens in blue, oxygens in red, and sulfur atoms in yellow. Hydrogen atoms are not shown. (**C**) Box plot comparing the range of the binding affinity experimental measurements used as references for the host-guest systems entering the SAMPL3 (purple), SAMPL4 (light blue), SAMPL5 (green), or SAMPL6 (yellow) challenges. The gray data points represent the measurements for the single host-guest entries. The the inter-quartile range and the median represented by the rectangular box were obtained by linear interpolation. The whiskers span the entire dynamic range of reported experiemntal measurements.

On the other hand, the median R^2^ improved with respect to the last round from 0.0 [0.0,0.8] to 0.5 [0.4,0.8]. Even in this case, we observe a slightly lower SAMPL6 median R^2^ when ignoring methods incorporating experimental data, but this is likely due not to the correction itself but to the fact that the top performing methods were generally submitted with and without correction, thus reducing the number of data points with high R^2^. Indeed, as already discussed, no positive effect on correlation was evident from the inclusion of a trained linear correction. The improvement is particularly evident when considering only free energy-based methodologies (e.g., alchemical and potential of mean force calculations). It should be pointed out out that the higher median R^2^ observed in SAMPL6 can, in principle, be explained not only by recent methodological advancements and the composition of the methods entering the challenge, but also by the particular set of assayed guests. While the first explanation is obviously the most desirable, the latter is a confounding factor when attempting to associate the results of the challenge to the progress of the community.

Since SOMD calculations entered the SAMPL5 challenge as well [19], we can compare directly the same statistics obtained by the method on the two guest sets to form an idea about the relative complexity of the two sets for free energy methods. To this end, we report in *Table 5* the uncertainties of the absolute statistics in terms of the mean and standard deviations of the bootstrap distributions instead of their 95-percentile confidence intervals to allow a direct comparison to those published in the SAMPL5 overview paper. The results of the SOMD methods applied to the OA/TEMOA-5 were submitted with a restraint and long-range dispersion correction, similarly to SOMD-C-nobuffer here, and without it, similarly to SOMD-A-nobuffer here. The two methods were referred as SOMD-3 and SOMD-1 respectively in the SAMPL5 overview. In both cases, the calculations used GAFF with AM1-BCC charges and TIP3P water molecules as well as a single flat-bottom restraint. The RMSE obtained by SOMD-C-nobuffer increased with respect to the statistic computed for SOMD-3 on OA/TEMOA-5 from 2.1 (2.1 ± 0.3) kcal/mol to 3.0 (3.0 ± 0.4) kcal/mol, where the number outside the parentheses is the statistic computed using all the data, and the numbers between parentheses are the mean and standard deviation of the bootstrap distribution. Incorporating experimental data into the prediction improved the error as SOMD-D-nobuffer obtained a RMSE of 1.6 (1.6 ± 0.3) kcal/mol. On the other hand, the Kendall correlation coefficient slightly increased on the SAMPL6 dataset from 0.4 (0.4 ± 0.2) to 0.7 (0.7 ± 0.4) while R^2^ remained more or less stationary from the already high value of 0.9 (0.7 ± 0.2) obtained on OA/TEMOA-5. Very similar observations can be made for SOMD-A-nobuffer and SOMD-1. While the improved τ correlation does not rule out the possibility of system-dependent effects on R^2^, it is unlikely for the difference between the median R^2^ of SAMPL5 and SAMPL6 (amounting to 0.76) to be entirely explained by the different set of guests, and the improvement is likely due, at least in part, to the different methodologies entering the challenge. In particular, SAMPL5 featured several free energy methods that scored near-zero R^2^ on the OA/TEMOA-5 set, affecting considerably the SAMPL5 median statistic. One of these methods is BEDAM, which used the OPLS-2005 [8, 58] force field and the implicit solvent model AGBNP2 [32], none of which entered the latest round of the challenge. However, the rest of these methods consist of double decoupling calculations carried out either with thermodynamic integration (TI) [60, 112] or HREX and BAR that employed CGenFF and TIP3P, which performed relatively well in SAMPL6 on OA/TEMOA. It should be noted that the TI and HREX/BAR methodologies in SAMPL5 made use of a Boresch-style restraint [18] harmonically constraining one distance, two angles, and three dihedrals. This is similar to the solution adopted in DDM-GAFF in SAMPL6, which also showed a relatively low R^2^ compared to the other free energy submissions in the same round of the challenge so it is natural to suspect that it may be particularly challenging to treat this class of host-guest systems with this type of restraint in alchemical calculations.

**Table 5.**
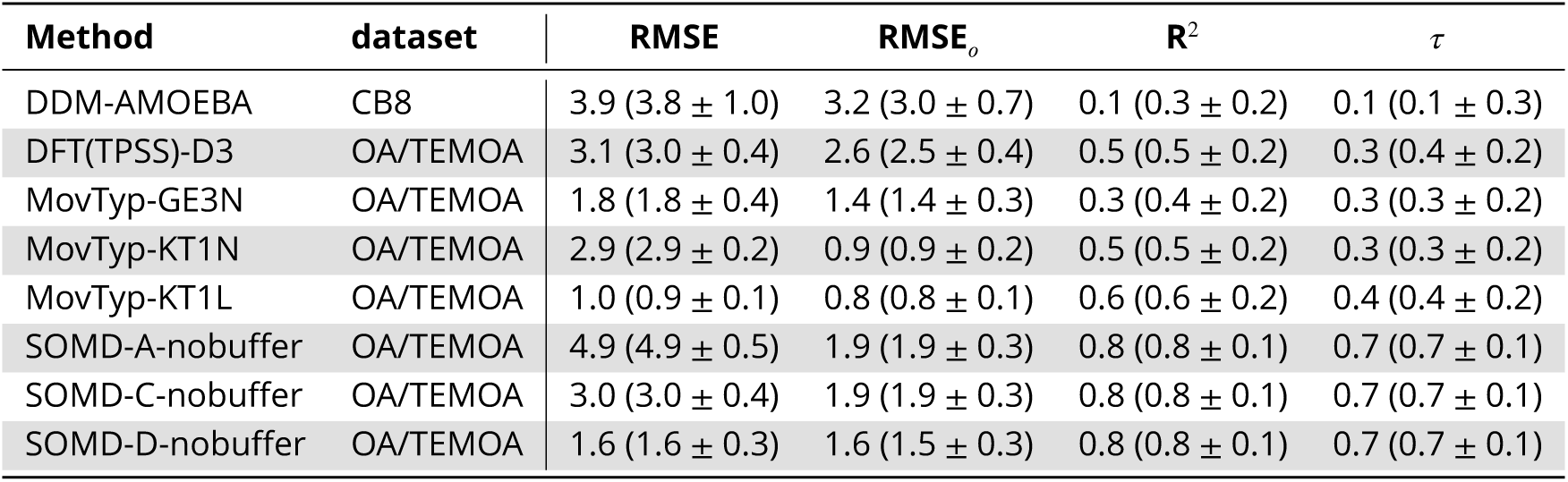
Offset statistics of the methods appearing in previous rounds of the SAMPL host-guest binding challenge. Root mean square error (RMSE), coefficient of determination (R^2^), Kendall correlation coefficient (*τ*), and offset root mean square error (RMSE_0_) computed by subtracting the mean signed error from the free energy predictions. Absolute and offset statistics for R^2^ and *τ* are identical, and they are thus reported only once. Absolute statistics are identical to those presented before, but, consistently with the format adopted in the SAMPL5 host-guest binding challenge overview paper, the are reported as mean ± standard deviation of the bootstrap distribution (between parentheses) instead of the 95-percentile confidence interval.

An improvement can also be observed for the movable type method, which was applied to the OA/TEMOA-5 set as well [10] using the KECSA1 and KECSA 2 potentials. These two submissions, identified with MovTyp-1 and MovTyp-2 respectively in the SAMPL5 overview paper, obtained similar statistics so we will use MovTyp-2 for the comparison. The SAMPL6 entry MovTyp-KT1N, which uses the KECSA energy model too, obtained a comparable RMSE of 2.9 (2.9 ± 0.2) kcal/mol against the 3.1 (2.9 ± 1.1) kcal/mol achieved by MovTyp-2 on OA/TEMOA-5, but, even in this case, the error becomes statistically distinguishable once the experimental-based correction is included (i.e., in MovTyp-KT1L), which decreases the RMSE to 1.0 kcal/mol. The correlation statistics generally compare favorably with respect to SAMPL5 with R^2^ moving from 0.0 (0.3 ± 0.3) to 0.5 (0.5 ± 0.2) and τ going from 0.1 (0.1 ± 0.3) to 0.3 (0.3 ± 0.2), although the uncertainties are too large to achieve statistical significance. Moreover, MovTyp-GE3N, which employs the more recently developed GARF energy model, obtained a better RMSE (1.8 (1.8 ± 0.4) kcal/mol) and comparable correlation statistics to MovTyp-KT1N.

Finally, it seems appropriate to compare the performance of DFT(TPSS)-D3 on OA/TEMOA to DFT/TPSS-c [21] in SAMPL5 and RRHO-551 [74] in SAMPL4 [86]. DFT(TPSS)-D3 an DFT/TPSS-c are very similar in that they both use the DFT-D3 approach to include dispersion correction, but while DFT(TPSS)-D3 generated an ensemble of configurations with MD, DFT/TPSS-c estimated the binding free energy from a single minimized structure. On the other hand, RRHO-551 does use MD for conformational sampling, but it employs DTF-D to correct for dispersion interactions, which was developed earlier than DFT-D3. As already mentioned, SAMPL4 featured a set of 9 OA guests [86], but only relative free energy predictions were submitted so absolute statistics are not available. Thus, in order to facilitate the comparison, we decided to report offset statistics for the subset of the SAMPL6 methods analyzed in this section in the same way they were computed in the previous two rounds of the challenge. The results are given in *Table 5*. The RMSE of the two models was relatively similar in SAMPL4 and SAMPL5: 5.8 ± 2.6 kcal/mol for RRHO-551 and 5.3 (5.2 ± 0.8) kcal/mol for DFT/TPSS-c, where the estimate for RRHO-551 does not include the mean of the statistic bootstrap distribution, which was not reported in the SAMPL4 overview paper. However, the SAMPL6 DFT(TPSS)-D3 calculations attained a lower error (2.6 (2.5 ± 0.4) kcal/mol) while maintaining a similar coefficient of determination of 0.5 (0.5 ± 0.2) against the 0.3 (0.4 ± 0.2) and 0.5 ± 0.2 of DFT/TPSS-c and RRHO-551 respectively.

#### The SAMPL6 CB8 system presents significant challenges to modern methodologies

A different perspective is offered by the history of the binding free energy predictions involving cucurbituril hosts. CB8 and the closely related CB7 appeared previously in SAMPL3 [87] together with an acyclic cucurbit[n]uril-type molecular container referred to as H1 [70]. Moreover, SAMPL5 featured another acyclic CB analogue called CBClip [128]. The 3D structures of the last two hosts are shown in *Figure 6*-B, while in *Figure 6*-A (bottom row), we show the distribution of RMSE and R^2^ computed from the binding free energy predictions submitted for SAMPL3 and SAMPL5 against these four hosts.

In general, both statistics appear to have deteriorated from SAMPL3 to SAMPL5. Even though H1 and CBClip are sufficiently different for system-dependent effects to reasonably dominate the overall performance, the most marked difference appears from the comparison of the SAMPL6 predictions to those submitted for CB7 and CB8 in SAMPL3, which achieved a much greater R^2^ in spite of the smaller dynamic range of the binding affinity measurements and none of which involved simulation-based methods. The explanation for this inequality is likely to be found in the complexity of the guest sets rather than a methodological regression as SAMPL3 featured only two relatively simple fragment-like binders while the latest round of the challenge included compounds of moderate size and/or complex stereochemistry (e.g., gallamine triethiodate, quinine).

That the CB8 guests in SAMPL6 were particularly challenging is corroborated by the comparison between the performance of DDM-AMOEBA and the results obtained by BAR-560, which also uses the double decoupling method and the AMOEBA polarizable force field, on the CB7 guests in SAMPL4 [14]. In this case as well, only offset statistics are available for comparison as SAMPL4 accepted exclusively relative free energy predictions. DDM-AMOEBA generally performed worse on the CB8 guest set featured in SAMPL6 with R^2^ decreasing from 0.6 ± 0.1 to 0.1 (0.3 ± 0.2) and RMSE increasing from 2.2 ± 0.4 to 3.2 (3.0 ± 0.7). While the CB8 guest set featured in SAMPL6 highlights the limits of current free energy methodologies, it also uncovers new learning opportunities that can be exploited to push the boundaries of the domain of applicability of these technologies.

Similarly to the OA/TEMOA guest set, simulation-based free energy methods display a higher median R^2^ than the global R^2^ computed from considering all the methods in the challenge, albeit a slightly higher RMSE as well. The pattern is consistent across the three rounds of the challenge, but the distributions of the statistics are too wide to draw statistically significant conclusions without collecting more data.

## Discussion

As in previous years, the SAMPL host-guest binding challenge has provided an opportunity for the computational chemistry community to focus on a common set of systems to assess the state-of-the-art practices and performance of current binding free energy calculation methodologies. The value of the blind challenge does not lie exclusively in the comparison and benchmarking of different methods, but also in its ability to highlight general areas of weakness in the field as a whole on which the community can focus. The latter aspect, in particular, risks to become of secondary importance in retrospective studies. Moreover, the consistent use of octa-acid and cucurbiturils since SAMPL3, which took place in 2011, give us the opportunity to make general observations over a longer time span.

### The variability in difficulty highlights the need to evaluate methodologies on the same systems

Several recurring themes have emerged from this and previous rounds of the challenge. Firstly, even for systems relatively simple as supramolecular host-guests, the performance of free energy methodologies and models can be heavily system-dependent. This is evident not only from the results of the same method applied to different guest sets, but also from the relative performance of the methods against different molecules. For example, most of the predictions employing GAFF obtained among the highest correlation statistics on the OA/TEMOA set while ranking among the lowest positions on the CB8 set. This stresses the importance of using the same set of systems when comparing multiple methodologies, which, without any coordination between groups, is a difficult task to carry out on a medium-large scale given the amount of expertise and resources necessary to perform this type of studies.

A useful dataset should be large enough to have the statistical power to resolve differences in performance, and diverse enough for the distribution of the binding affinity to approximate the distribution of the population of interest and reflect how the method would perform on new data. At the same time, however, correlation statistics tend to increase with the dynamic range spanned by the data, and some methods, such as relative free energy calculations, often impose practical limits to the structural differences between compounds. For example, RFEC-GAFF and RFEC-QMMM submitted predictions only for the OA/TEMOA set, where the similarities between the guests are more prominent. These contrasting requirements, together with practical problems connected to the availability of experimental data and resources, make crafting an appropriate dataset a very challenging task.

### Force field accuracy is a dominant limiting factor for modeling affinity

A second consideration surfacing from previous SAMPL rounds as well is the tendency of classical methods to overestimate the binding affinities. Since the results of the related SAMPLing challenge support the claim that convergence for this class of systems is achievable [51], and considering that the RMSE has not improved significantly across rounds of the challenge, this seem to suggest that an investment of resources into improving the empirical parameters of force fields and solvent models could have a dramatic impact. It should be noted that, while these systems do not put to the test protein parameters, they rely on general force fields that are routinely used in drug and small molecule design.

### Other missing chemical details may also be major limiting factors

However, the problem of missing details of the chemical environment such as salts and alternative protomers cannot be ruled out as a major determinant of predictive accuracy. Explicitly modeling the buffer salt concentrations in the SOMD-C predictions reduced the RMSE from 7.9 to 5.1 kcal/mol for two sets of simulations otherwise identical, and, curiously, it had the opposite effect of increasing the error statistics on the OA/TEMOA set. Despite the sensitivity of the free energy prediction to the presence of ions, a lack of standard best practices emerges from the challenge entries. Many participants decided to add only neutralizing counterions or use a uniform neutralizing charge, and others did not include information about how the buffer was modeled in the submitted method sections, which possibly reflects a generally minor role currently played by this particular aspect of the decision-making process during the modeling step in comparison to other elements (e.g., charges, force field parameters, water model).

### Even at extreme pH, protonation state effects may still contribute

The possible influence of multiple accessible protonation states of the guest compounds on the binding free energy was left unexplored during the challenge, mirroring the widespread tendency in the free energy literature to neglect its effect, and participants largely used the most likely protonation states predicted by Epik that were provided in the input `mol2` and `sdf` files. However, the p*K*_*a*_ free energy penalties estimated by Epik for the second most probable protonation state of the CB8 guests in water at experimental pH (*Table 6*), which is obtained in all cases by the deprotonation of the charged nitrogen atoms as given in *Figure 1*, suggest that for several guests, and in particular for CB8-G3 and CB8-G11, the deprotonated state is accessible by paying a cost of a few *k*_B_*T* (where *k*_*B*_ is Boltzmann’s constant and *T* is the absolute temperature), and a change in relative populations between the end states driven by the hydrophobic binding cavity may have a non-negligible effect on the binding affinity. Furthermore, even if the probability of having the carboxyl group of the octa-acid guests protonated at pH 11.7 is usually neglected, a previous study performed for SAMPL5 showed that modeling changes in protonation state populations upon binding resulted in improved predictive performance for a set of OA and TEMOA guests that, similarly to the latest round of the challenge, included several carboxylic acids and was measured at a similar buffer pH [118]. Experimentally, net proton gain or loss during complexation could straightforwardly be assessed for highly soluble host-guest systems via isothermal titration calorimetry (ITC) in buffers with the same pH but different ionization heats for proton loss from solvent [7], a technique that has been used for protein-ligand systems [24, 25, 91, 111]. Similarly to buffer salts, there are few established practices in the community to treat multiple protonation states in free energy calculations [66], and further development and testing of force fields and solvent models with the goal of improving accuracy to experiments should consider these issues as ignoring them during the fitting procedure could push the error caused by missing essential chemicals (e.g., ions, protonation and tautomeric states) to other force field parameters with the risk of decreasing the transferability of the model.

**Table 6.**
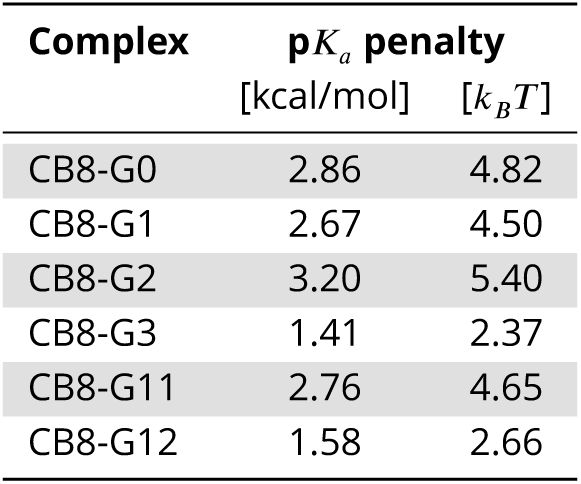
p*K*_*a*_ free energy penalties predicted by Epik for the second most likely protonation state of the CB8 guests. In all cases, the second most probable protonation state predicted by Epik can be obtained by removing the nitrogen proton of the dominant state. The estimated free energy penalties to access the deprotonated state are reported in kcal/mol and units of *k*_*B*_*T*, where *k*_*B*_ is the Boltzmann’s constant and *T* is the absolute temperature, taken to be 298 K (i.e., the temperature at experimental conditions). For all the other compounds, including the octa-acid guests, Epik was not able to find a second protonation state within a tolerance of 3 pH units.

| Complex | $pK_a$ penalty | |
| --- | --- | --- |
| | [kcal/mol] | $[k_B T]$ |
| CB8-G0 | 2.86 | 4.82 |
| CB8-G1 | 2.67 | 4.50 |
| CB8-G2 | 3.20 | 5.40 |
| CB8-G3 | 1.41 | 2.37 |
| CB8-G11 | 2.76 | 4.65 |
| CB8-G12 | 1.58 | 2.66 |

### Linear corrections ft to prior experimental measurements do not improve predictive utility

The experimental-based correction adopted by several groups introduces a new theme in the challenge which pertains to strategies that can be used to inject previous knowledge into molecular simulations. Force field parameters are in principle capable of incorporating experimental data, but an update of the model driven by binding free energy measurements or other ensemble observables is doubtlessly challenging and may involve calculations as expensive as the production calculations so this is normally not routinely viable, although previous studies indicated the validity and feasibility of such an approach [125, 126]. Other schemes that emerged in particular from the field of crystallographic structural refinement avoid modifying the force field parameters and instead add one or more biasing terms to the simulation to replicate experimental measurements that the underlying force field cannot reproduce [16, 123]. The simple linear corrections used independently by various participants in this round of the challenge had a positive impact on the error, but a very small effect in terms of correlation, which is often of central importance in the context of molecular design. However, the simplicity of its application, which is confined entirely to the post-processing step, was such that the participants were able to submit multiple entries with and without the correction.

### Outlook for future SAMPL host-guest challenges

The SAMPL roadmap [77] outlines a proposal for subsequent host-guest challenges for SAMPL7-10. While the future of these blind exercises is uncertain given the absence of a sustainable funding source, we briefly review the likely future design of these host-guest challenges below.

In one line of exploration ([77], section 2.2), SAMPL7 proposes to explore variants of Gibb deep cavity cavitands (related to OA/TEMOA) in which carboxylate substitutent locations are modified, comparing multiple host variants against a set of guests to explore how well affinities and selectivities could be predicted.

SAMPL8 would provide a second iteration of this experiment with novel guests and a trimethylammonium-substituted hostvariantto assess how algorithmic improvements from the first round could lead to improved performance. SAMPL9-10 would consider the effect of common biologically relevant salts, comparing the effects of NaCl and NaI on various host variants, while SAMPL11 would consider the effects of cosolvents that might compete for the binding site or modulate the strength of the hydrophobic effect.

In another line of exploration ([77], section 2.1), SAMPL7-11 are also proposed to feature cucubituril variants, including methylated forms of CB8, glcoyuracil hexamer, and acyclic forms of CB[n]-type receptors. By comparing the constrained cyclic and less constrained acyclic forms of CB[n] hosts, the accuracy with which participants can model the energetics of receptor flexibility and receptor desolvation can be probed. SAMPL8-9 also plans to feature small molecule guests with pKa values between 3.8–7.4, which brings the possibility that host binding can induce substantial shifts in protonation state.

Finally, recent work by one of the authors has demonstrated how a library of monosubstituted β-cyclodextrin analogues can be generated via a simple chemical route [59]. This strategy could ultimately lead to the attachment of chemical groups that resemble biopolymer residues, such as amino or nucleic acids, allowing interactions between small druglike molecules and biopolymer-like functional groups to be probed without the multifold challenges that protein-ligand interactions present. While development of this system is still ongoing, it is likely to make an appearance in upcoming SAMPL host-guest challenges.

## Code and data availability

- Input files and setup scripts: https://github.com/MobleyLab/SAMPL6/tree/master/host_guest/
- Analysis scripts: https://github.com/MobleyLab/SAMPL6/tree/master/host_guest/Analysis/Scripts/
- Analysis results: https://github.com/MobleyLab/SAMPL6/tree/master/host_guest/Analysis/Accuracy
- Participants’ submissions: https://github.com/MobleyLab/SAMPL6/tree/master/host_guest/Analysis/Submissions

## Author Contributions

Conceptualization, AR, JDC, DLM; Methodology, AR, JDC, DLM; Software, AR; Formal Analysis, AR, JDC; Investigation, AR, QY, SM, MS, JNM; Resources, JDC, BCG, LI, MWC, MKG, DLM; Data Curation, AR, MWC; Writing-Original Draft, AR, JDC; Writing - Review and Editing, AR, JDC, DLM, MKG, LI, BCG, SM; Visualization, AR, SM; Supervision, JDC, DLM; Project Administration, AR,JDC, DLM; Funding Acquisition, JDC, DLM, MKG, BCG, LI.

## Acknowledgments

AR and JDC acknowledge support from the Sloan Kettering Institute. JDC acknowledges support from NIH grant P30CA008748. JDC, AR, and DLM gratefully acknowledge support from NIH grant R01GM124270 supporting SAMPL blind challenges. AR acknowledges partial support from the Tri-Institutional Program in Computational Biology and Medicine. LI thanks the National Science Foundation for supporting (CHE-1404911) the participation in SAMPL6. DLM appreciates financial support from the National Institutes of Health (1R01GM108889-01), the National Science Foundation (CHE 1352608). AR and JDC are grateful to OpenEye Scientific for providing a free academic software license for use in this work. We thank four anonymous reviewers, whose comments helped us improve the manuscript. The content is solely the responsibility of the authors and does not necessarily represent the official views of the National Institutes of Health.

## List of abbreviations

AM1-BCC: Austin model 1 bond charge correction [54, 55]
AMOEBA: atomic multipole optimized energetics for biomolecular simulation [96]
B3LYP: Becke 3-parameter Lee-Yang-Parr exchange-correlation functional [13]
B3PW91: Becke 3-parameter Perdew-Wang 91 exchange-correlation functional [13]
CGenFF: CHARMM generalized force field [120]
COSMO-RS: conductor-like screening model for real solvents [61]
DDM: double decoupling method [39]
DFT-D3: density functional theory with the D3 dispersion corrections [42]
FM: Force Matching [28]
FSDAM: Fast switching double annihilation method [92, 97]
GAFF: generalized AMBER force field [121]
HREX: Hamiltonian replica exchange [113]
KECSA: knowledge-based and empirical combined scoring algorithm [129]
KMTISM: KECSA-Movable Type Implicit Solvation Model [131]
MD: molecular dynamics
MMPBSA: molecular mechanics Poisson Boltzmann/solvent accessible surface area [110]
MovTyp: Movable Type method [130]
OPLS3: optimized potential for liquid simulations [45]
PBSA: Poisson-Boltzmann surface area [106]
PM6-DH+: PM6 semiempirical method with dispersion and hydrogen bonding corrections [64,100]
RESP: restrained electrostatic potential [12]
REST: replica exchange with solute torsional tempering [68, 71]
RFEC: relative free energy calculation
QM/MM: mixed quantum mechanics and molecular mechanics
SOMD: double annihilation or decoupling method performed with Sire/OpenMM6.3 software [27, Woods et al.]
SQM: semi-empirical quantum mechanics
TIP3P: transferable interaction potential three-point [57]
TPSS: Tao, Perdew, Staroverov, and Scuseria exchange functional [116]
US: umbrella sampling [119]
VSGB2.1: VSGB2.0 solvation model reft to OPLS2.1/3/3e [67]

## Supplementary figures

**Appendix 0 Figure 7.**
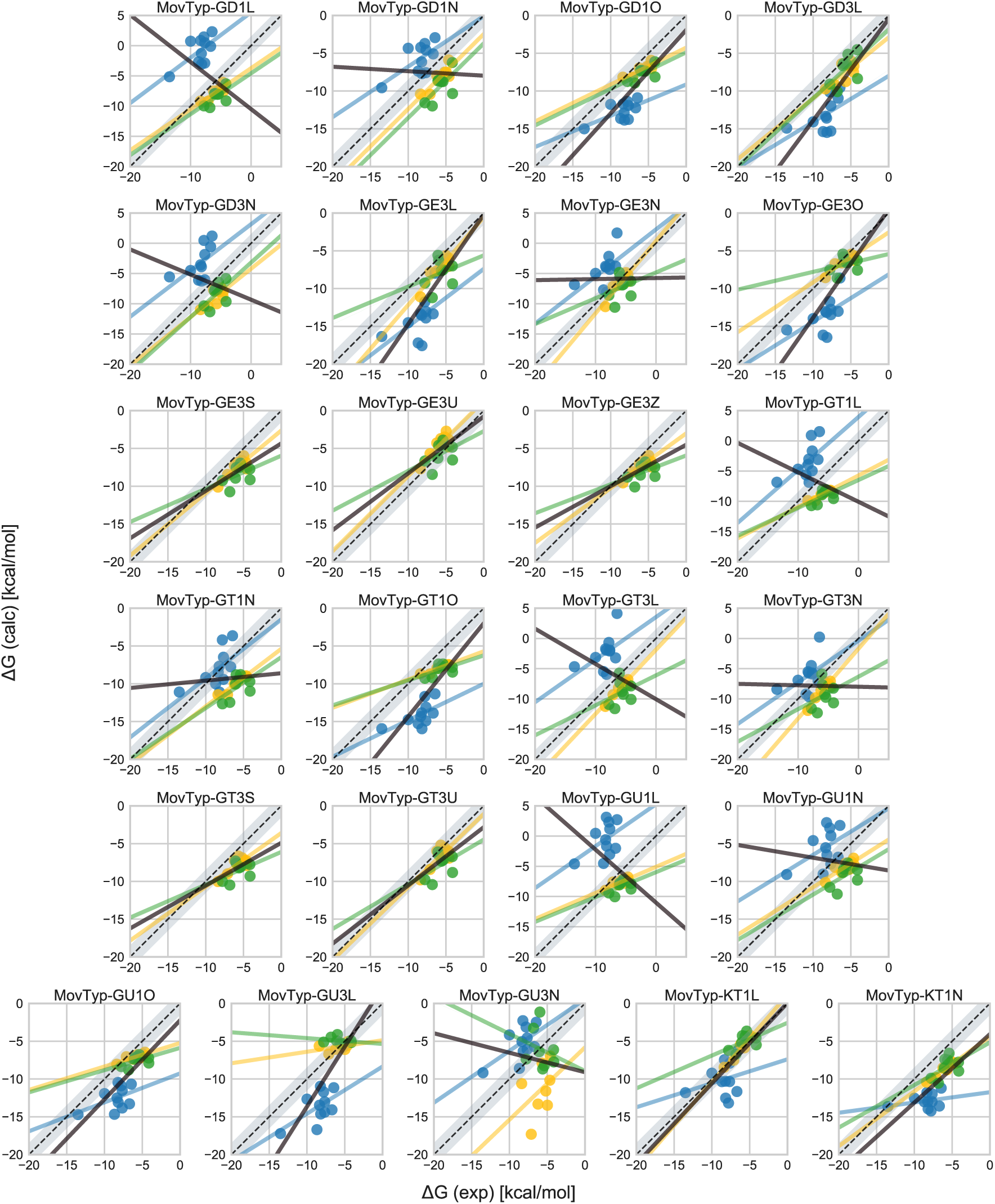
Free energy correlation plots of all movable type submissions. The four-letter suffix of each movable type submission is to be interpreted as following: first letter indicates the force field (G: GARF; K: KECSA), the third letter input structures (D: final frame of MD sampling; E: ensemble of structures from MD sampling; T: lowest energy structure during movable type scoring; U: lowest energy structure obtained during the sampling in US-GAFF), the third letter is the number of states (1: only the complex is considered, 3: includes also host and guest in solution), and the fourth letter the type of experimental correction (N: no correction; L: linear correction trained a single dataset including OA, TEMOA, and CB8; O: offset correction trained a single dataset including OA, TEMOA, and CB8; U: linear correction trained on a set excluding CB8 guests; S: two different linear corrections for OAand TEMOA predictions trained on two separated sets including either OA or TEMOA measurements; Z: same as S but with only offset term).

**Appendix 0 Figure 8.**
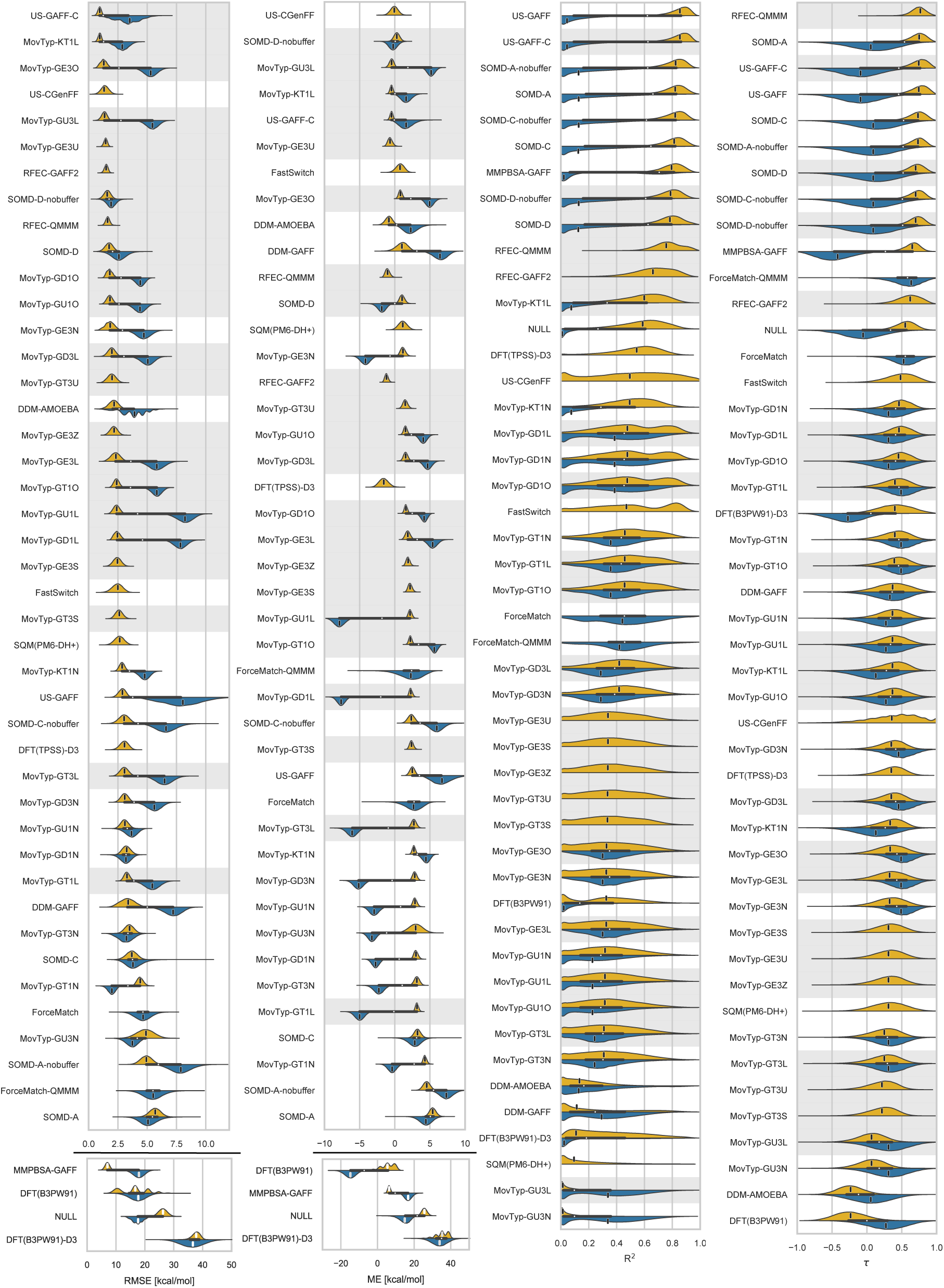
Bootstrap distributions including all the movable type submissions.

**Appendix 0 Figure 9.**
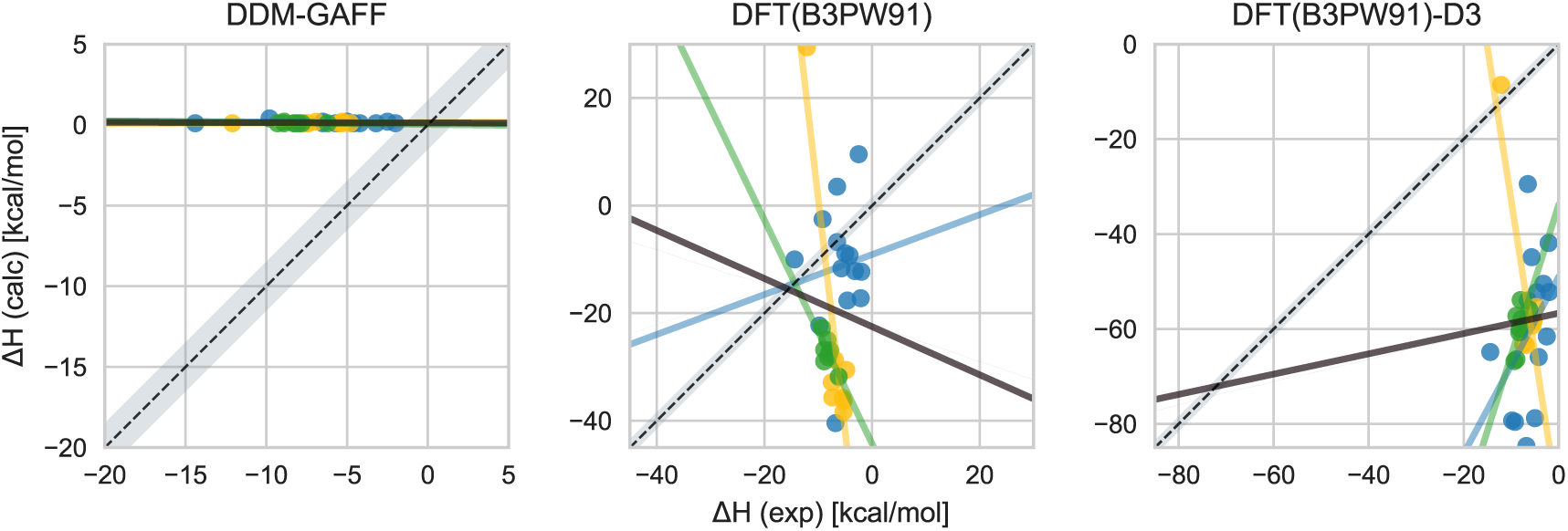
Enthalpy correlation plots obtained by the methods on the three host-guest sets. Scatter plots showing the experimental measurements of the host-guest binding enthalpies (horizontal axis) against the methods’ predictions on the OA (yellow), TEMOA (green), and CB8 (blue) guest sets with the respective regression lines of the same color. The solid black line is the regression line obtained by using all the data points. The gray shaded area represent the points within 1.5 kcal/mol from the diagonal (dashed black line).

## Footnotes

1 Technically, this is an aZne transformation in the general case since *b* ≠ 0 for some of the corrections employed by participants, but we will refer to it as *linear* here.

